# Predicting Tumor Response to Drugs based on Gene-Expression Biomarkers of Sensitivity Learned from Cancer Cell Lines

**DOI:** 10.1101/2020.07.03.180620

**Authors:** Yuanyuan Li, David M. Umbach, Juno Krahn, Igor Shats, Xiaoling Li, Leping Li

**Affiliations:** Biostatistics and Computational Biology Branch, National Institute of Environmental Health Sciences, Research Triangle Park, NC 27709, USA; Genome Integrity & Structural Biology Laboratory, National Institute of Environmental Health Sciences, Research Triangle Park, NC 27709, USA; Signal Transduction Laboratory, National Institute of Environmental Health Sciences, Research Triangle Park, NC 27709, USA

**Author notes:** Correspondence (L.L.).

## Abstract

Human cancer cell line profiling and drug sensitivity studies provide valuable information about the therapeutic potential of drugs and their possible mechanisms of action. The goal of those studies is to translate the findings from *in vitro* studies of cancer cell lines into *in vivo* therapeutic relevance and, eventually, patients’ care. Tremendous progress has been made. In this work, we built predictive models for 453 drugs using data on gene expression and drug sensitivity (IC_50_) from cancer cell lines. We identified many known drug-gene interactions and uncovered several potentially novel drug-gene associations. Importantly, we further applied these predictive models to ∼17,000 bulk RNA-seq samples from The Cancer Genome Atlas (TCGA) and the Genotype-Tissue Expression (GTEx) database to predict drug sensitivity for both normal and tumor tissues. We created a web site for users to visualize and download our predicted data (https://edelgene.niehs.nih.gov/cancerRxTissue). Using trametinib as an example, we showed that our approach can faithfully recapitulate the known tumor specificity of the drug. We further demonstrated that our approach can predict drugs that 1) are tumor-type specific; 2) elicit higher sensitivity from tumor compared to corresponding normal tissue; 3) elicit differential sensitivity across breast cancer subtypes. If validated, our predictions could have clinical relevance for patients’ care.

## INTRODUCTION

Studies that characterize human cancer cell lines and evaluate their sensitivity to drugs provide valuable information about the therapeutic potential and the possible mechanisms of action of those drugs. Those studies allow the identification of genomic features that are predictive of drug responses and make it possible to relate findings from cell lines to tissue samples and, ultimately, to translate laboratory results into patients’ care.

The Genomics of Drug Sensitivity in Cancer (GDSC) Project has assayed the sensitivity of 987 cancer cell lines to 320 compounds in their phase 1 (GDSC1) assay and of an additional 809 cancer cell lines to 175 compounds (some of which were included in the GDSC1 assay) in their phase 2 (GDSC2) assay (Garnett et al., 2012; Iorio et al., 2016; Yang et al., 2013). The sensitivity of each cancer cell line to the drugs was represented as an IC_50_ value (the concentration at which a cell line exhibited an absolute inhibition in growth of 50%; lower IC_50_ implies higher sensitivity). GDSC also quantified the basal level gene expression of many of the cancer cell lines using microarray (Garnett et al., 2012). Concomitantly, other consortia such as the CCLE (cancer cell line encyclopedia) also profiled genome-wide gene expression of many of the cancer cell lines using RNA-seq (Barretina et al., 2012; Ghandi et al., 2019). Additional genomic features such as somatic mutation and copy number variation, DNA methylation, epigenetic modifications, microRNA expression, and protein expression were also characterized by CCLE and others (Barretina et al., 2012; Ghandi et al., 2019). The Cancer Therapeutics Response Portal (CTRP) project profiled the sensitivity of 860 cancer cell lines to 481 small molecules (Rees et al., 2016; Seashore-Ludlow et al., 2015). The National Cancer Institute (NCI) has carried out a screening assay for a large number of small molecule compounds to detect potential anticancer activity using a group of 60 human cancer cell lines (NCI60) (Reinhold et al., 2012). Recently, the transcriptomes of the NCI-60 cancer cell lines were also analyzed using RNA-seq (Reinhold et al., 2019). Those resources make it possible to associate sensitivity of cancer cells to different drugs with genomic information on the cells, thereby, facilitating the discovery of molecular biomarkers of sensitivity and the identification genomic and genetic features that are predictive of cell sensitivity (Barretina et al., 2012; Rajapakse et al., 2018; Reinhold et al., 2012; Reinhold et al., 2019).

GDSC applied various statistical and computational methods including elastic net regression and machine learning algorithms to identify multiple interacting genomic features influencing each cell line’s sensitivity to drugs. These analyses identified many interactions between cancer gene mutations and specific drugs (Yang et al., 2013). For example, cancer cell lines with mutations in the BRAF genes are significantly more sensitive to PLX4730, a BRAF-inhibitor, than those with wild-type BRAF (Yang et al., 2013). Additional computational methods for predicting sensitivity to drugs using gene expression data from cancer cell lines have been developed, for example (Azuaje et al., 2018; Nguyen et al., 2016; Suphavilai et al., 2018; Wei et al., 2019). Several publications have recent efforts in this area (Azuaje, 2017; Guan et al., 2019; Guvenc Paltun et al., 2019; Reinhold et al., 2015).

In addition to efforts directed at understanding the relationship between drug sensitivity and the genomic and genetic characteristics of cancer cell lines, major efforts have been put into relating the findings from *in vitro* studies of cancer cell lines to *in vivo* relevance. For example, Iorio et al. (Iorio et al., 2016) carried out a comprehensive characterization of genomic alternations including somatic mutations, copy number alterations, and DNA methylation in 11,289 tumors and 1,001 cancer cell lines. Tumor sensitivity to 265 drugs was predicted using corresponding sensitivity data from cancer cell lines by mapping cancer-driven alterations *-* the cancer functional events (CFEs) *-* in the tumors to cancer cell lines (Iorio et al., 2016). The authors identified single CFEs or combinations of them as markers of response and used a deep learning method to identify associations between drug molecular descriptors and mutational fingerprints in cancer cell lines. They subsequently used such associations to predict the potential of repurposing FDA approved drugs for cancer treatment (Chang et al., 2018). Similarly, DeepDR considered genomic profiles of both cancer cell lines and tumors to predict tumor sensitivity to drugs using a deep neural network (Chiu et al., 2019).

Those genomic-based approaches uncovered oncogenic alterations that are susceptible to anti-cancer drugs, thereby helping to identify treatment options that are tethered to specific genetic aberrations. Conversely, other approaches relate cell-line findings to tumors based on transcriptome data. For example, using expression and drug sensitivity data from cancer cell lines, Geeleher et al. (Geeleher et al., 2014; Geeleher et al., 2017) developed gene expression-based models to predict sensitivity to drugs; they subsequently applied the models to gene-expression data from TCGA tumor samples to impute sensitivity of the tumor samples to 138 drugs.

Similarly, we sought to identify gene expression signatures from cancer cell lines that can predict their sensitivity to drugs; and, subsequently, we used those signatures to predict the sensitivity of normal and tumor tissue to the drugs. Our work, however, differs from the previous work in several ways: a) our analysis is more comprehensive by including the latest drug sensitivity data from GDSC2 for 453 drugs; b) our work emphasizes identification of putative biomarkers of sensitivity to drugs and potential therapeutic options for cancer subpopulations; and c) we also predict toxicity of drugs to normal tissues using transcriptomic data from normal human tissues available from both The Cancer Genome Atlas (TCGA) and The Genotype-Tissue Expression (GTEx) project.

We identified many known drug-gene interactions and uncovered several potentially novel drug-gene associations. We predicted that OSI-027 (mTOR inhibitor) is a breast cancer specific drug with high specificity for the Her2-positive subtype breast tumors. Our analysis also suggests that *MULC1* expression is a surrogate marker for tumor response to OSI-027. Our analysis rediscovered the interaction between bleomycin and ACE (angiotensin I converting enzyme) (Day et al., 2001). We also predicted that other drugs are potentially specific for cancer (sub)types. If validated, our predictions could have clinical relevance for patients’ care.

## RESULTS

Using cancer cell line gene expression data and cancer cell line drug sensitivity data, we built predictive models that were subsequently used to impute/predict tissue drug sensitivity using gene expression data for the tissues (Figure 1). Details are provided in Methods.

**Figure 1.**
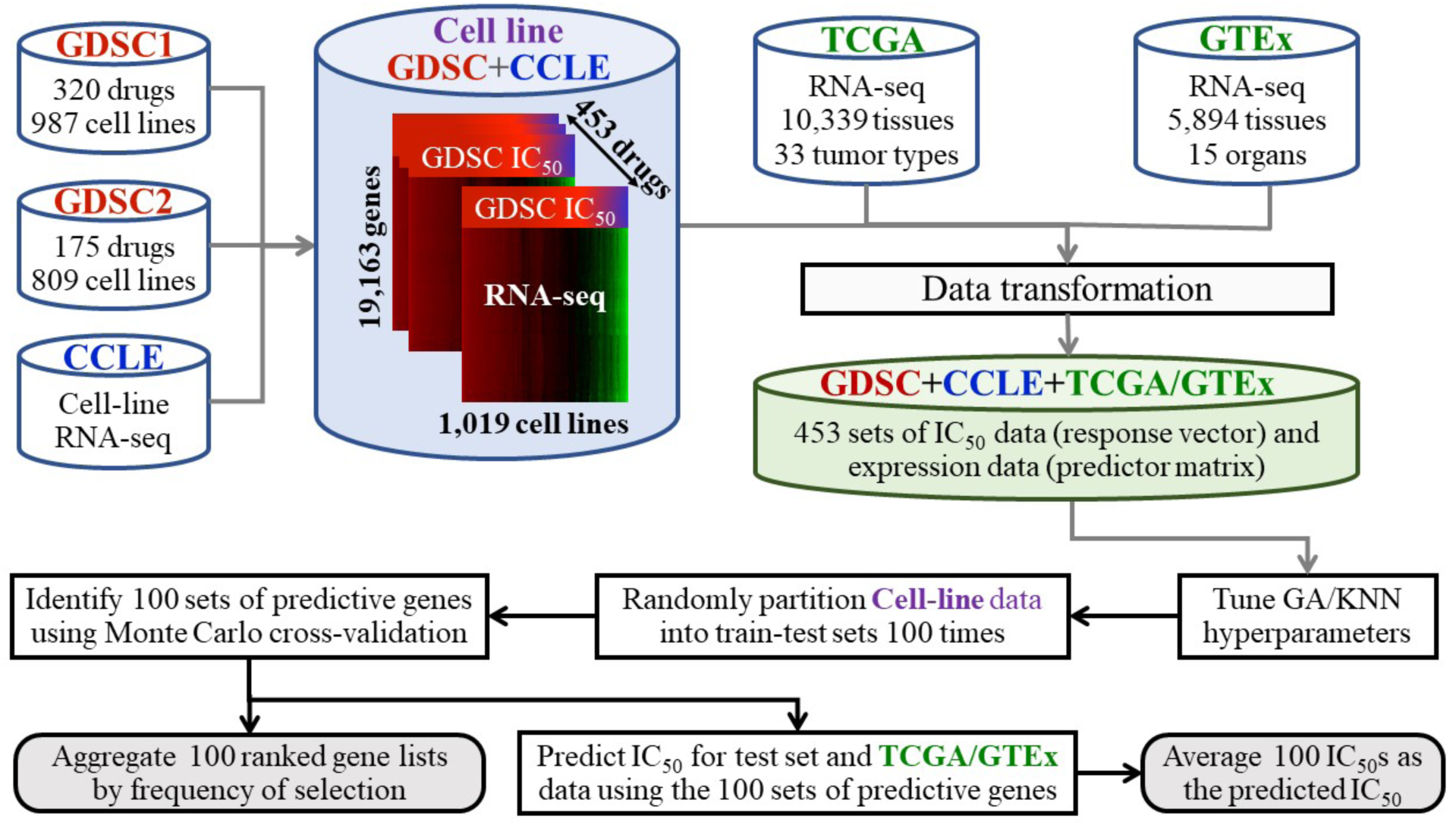
Schematic diagram of the work flow. First, GDSC cancer cell line drug sensitivity data, CCLE cancer cell gene expression data and TCGA/GTEx tissue gene expression data are combined and transformed. The CCLE gene expression data and GDSC drug sensitivity data (collectively referred to as the cell-line data) were used to build predictive models that were subsequently used to predict/impute the tissue drug sensitivity for the TCGA and GTEx samples. Broadly, for each drug, we divided the cell-line data into a training and testing set. We aimed to identify a 30-gene set whose gene expression levels are most predictive of the IC_50_ values of the drug for the samples in the testing set. The resulting model (a 30-gene set) was subsequently used to predict the IC_50_ value of the TCGA/GTEx samples. This process was repeated 100 times independently. The predicted IC_50_ values from the 100 runs were then averaged and taken as the predicted IC_50_ value of the drug for the samples. For details, see Methods.

### Training and Testing Performance for the Cell-Line Data

First, we divided the cancer cell-line data into a training and testing set. We predicted the IC_50_ values of the cancer cell lines in the testing set for each of the 453 drugs. We computed both the Pearson (*ρ*_*P*_) and Spearman (*ρ*_*S*_) correlation coefficients between the observed and predicted IC_50_ values for the samples in the testing set. The median *ρ*_*P*_ and *ρ*_*S*_ coefficients between the observed and predicted IC_50_ values were 0.466 and 0.437 (Table S1), indicating that the basal transcriptomes of the cancer cell lines can reasonably predict the sensitivity (IC_50_s) of the cell lines to most of the drugs. Of the 453 drugs, 272 (60%) had both *ρ*_*P*_ and *ρ*_*S*_ testing-set correlations ≥ 0.4. We refer to those drugs as predictable drugs.

Interestingly, for 34 (7.5%) of the drugs, the cancer cell lines’ transcriptomes had little or no predictive power for the cell lines’ sensitivities to the drug (either *ρ*_*P*_ or *ρ*_*S*_ testing-set correlation coefficients ≤ 0.25). Moreover, we also confirmed that other transcriptomic data such as microRNA expression, DNA methylation, and protein expression (from reverse phase protein array) from CCLE and GDSC (Barretina et al., 2012; Ghandi et al., 2019) also had little or no predictive power for those drugs (data not shown). It is unclear why the transcriptomes of cancer cell lines failed to predict their sensitivity to those drugs. Some of 34 drugs had fewer than 100 samples with both gene expression and IC_50_ data and the lack of data may have contributed to those drugs’ poor prediction performance; for most of the others, however, data availability was not an issue.

For the remaining analyses, we focus on the top 272 predictable drugs – those having the highest testing-set correlations between the observed and predicted IC_50_ values (both *ρ*_*P*_ ≥ 0.4 and *ρ*_*S*_ ≥ 0.4). The top 10 predictable drugs (Table S2) appear to have diverse mechanisms of action.

### Top-ranked Genes Predictive of Drug Sensitivity

For each of the top 272 predictable drugs, we counted how many times each gene was selected into the 100 sets of the *d* (*d*=30) predictive genes. For a transcriptome of 19,163 genes, a gene is expected by chance to be selected only 0.155 times [(30/19163) × 100] into 100 sets of 30 genes. We observed that many genes were being selected at frequencies more than 100 times above that expected by chance. The most frequently selected genes for each drug are potentially informative about that drug’s mechanism of action as well as about a cancer cell line’s sensitivity to the drugs. For some other drugs, multiple genes were selected with lower but distinctly higher-than-random frequencies, suggesting that multiple genes together are necessary for predicting cell-line sensitivity for those drugs. Many of the drug-gene interactions were also identified by others (Barretina et al., 2012; Garnett et al., 2012; Iorio et al., 2016). Among the predictable drugs, the number of genes selected into more than 20 of 100 predictive gene sets (i.e., > 100-fold above chance) ranged from 1 to 17 (Table S3).

*C19orf33* (chromosome 19 open reading frame 33) was among the most frequently included predictive genes for the largest number of drugs, appearing in more than 20% of the predictive gene sets for 17 drugs (Table S3). The expression level of *C19orf33* in cancer cell lines was positively correlated (*ρ*_*S*_ > 0.3) with the IC_50_ values of more than 100 drugs for those cell lines (Table S4), suggesting that higher expression of *C19orf33* in cancer cell lines was positively associated with higher resistance of the cancer cell lines to the drugs. No other genes were correlated with the IC_50_ values of as many drugs as was *C19orf33*. Most of the positively correlated drugs are DNA synthesis inhibitors, microtubule assembly inhibitors, or cell cycle inhibitors. Interestingly, *C19orf33* expression in cancer cell lines showed a negative correlation with the IC_50_ values of the kinase (MEK, ERK, SRC) inhibitors for the cell lines (Table S4), suggesting that cancer cell lines with higher *C19orf33* expression are more sensitive to kinase inhibitors than those with lower *C19orf33* expression. *C19orf33* encodes two transcript variants (Immortalization up-regulated protein 1 and 2: IMUP-1 and IMUP-2); both were discovered and characterized in immortalized cells as being upregulated compared to senescent cells (Kim et al., 2000). IMUP-1 and IMUP-2 are more frequently expressed in cancer cells compared to normal tissues (Kim et al., 2000; Kim et al., 2008; Uchiyama et al., 2007). Overexpression of IMUP-1 and IMPU-2 in normal fibroblasts induces neoplastic transformation (Ryoo et al., 2006). Our data suggested that *C19orf33* expression may be a general biomarker for the sensitivity of cancer cell lines to many chemotherapeutic agents.

For 14 drugs of diverse mechanisms of action, *ABCB1* appeared in more than 20% of the predictive gene sets. *ABCB1* encodes ATP binding cassette subfamily B member 1, a member of the superfamily of ATP-binding cassette (ABC) transporters. ABCB1, also commonly known as MDR1, is an ATP-dependent drug efflux pump for xenobiotic compounds with broad substrate specificity (Hodges et al., 2011). Expression of *ABCB1* is responsible for decreased drug accumulation in multidrug-resistant cells and often mediates the development of resistance to anticancer drugs (Chen and Sikic, 2012; Hodges et al., 2011; Robey et al., 2018).

*SLFN11* was another frequently selected gene for multiple drugs (Table S3). *SLFN11* encodes the Schlafen family member 11 and is a putative DNA/RNA helicase. In the NCI-60 cell lines, *SLFN11* expression is positively correlated with cell-line sensitivity not only to topoisomerase inhibitors but also to DNA alkylating and synthesis inhibitors (Zoppoli et al., 2012). In clinical trials, tumors expressing SLFN11 treated with temozolomide exhibit enhanced cell death following DNA damage (Pietanza et al., 2018). *SLFN11* expression in the CCLE cancer cell lines was inversely correlated with the IC_50_ values of the drugs for which *SLFN11* was the most frequently selected gene (Figure S2), indicating that cancer cell lines with higher *SLFN11* expression were more sensitive to those drugs. Those results suggest that *SLFN11* may also be a general biomarker for the effectiveness of chemotherapeutic agents.

#### Drugs whose top-ranked predictive genes matched the drug targets

Many of the most frequently selected genes were also known targets of the drugs (Table S3). The drug nutlin-3a inhibits the interaction between p53 and MDM2, leading to activation of the p53 pathway (Vassilev et al., 2004). Gratifyingly, *MDM2* was selected in nearly 100% of gene sets predicting sensitivity to nutlin-3a. Other known p53 known target genes (*CDKN1A* and *RPL22L1*) were also selected in nearly 90% of those predictive gene sets, suggesting that high expression of these genes identifies tumors with a wild-type p53, and thus responsive to a further p53 activation by nutlin-3a. The drugs PD173074 and AZD4547 are two potent fibroblast growth factor receptor (FGFR) inhibitors (Gudernova et al., 2016); *FGFR2* was the most frequently selected gene for predicting sensitivity to these two drugs. Venetoclax is known to target BLC2 protein (Anderson et al., 2016); *BCL2* was selected in almost 100% of the predictive gene sets for sensitivity to venetoclax. Tanespimycin (also known as 17-AAG) is a HSP90 inhibitor, and *NQO1* expression is inversely correlated with 17-AAG IC_50_s in cancer cell lines (Gaspar et al., 2009). Similarly, we found that *NQO1* was selected in 100% in the predicted gene sets for tanespimycin sensitivity. *ADK* (adenosine kinase) was selected in 100% of gene sets predicting sensitivity to AICAR. AICAR is an analog of adenosine monophosphate (AMP) that is capable of stimulating AMP-dependent protein kinase (AMPK) activity (Merrill et al., 1997). AICAR prevents the production of the enzymes adenosine kinase (ADK) and adenosine deaminase (ADA) (Boison, 2013).

*SPRY2* (sprouty RTK signaling antagonist 2) was among the most frequently selected genes for predicting sensitivity to <u>all</u> six MEK1/2 inhibitors in GDSC datasets (CI-1040, PD0325901, refametinib, SCH772984, selumetinib, and trametinib) (Table S3). Sprouty specifically inhibits activation of MAPK/ERK in response to a wide range of trophic growth factors (Gross et al., 2001; Masoumi-Moghaddam et al., 2014). *SPRY2* expression in cancer cell lines was inversely correlated with the IC_50_ values of all MEK inhibitors for the cancer cell lines (Figure S3) with *ρ*_*S*_ ranging from -0.24 to -0.43 (all *p*-values < 4E-08), indicating that cancer cell lines with higher *SPRY2* expression are more sensitive to the MEK inhibitors. The other most frequently selected genes for MEK1/2, BRAF, and ERK1/2 kinase inhibitors were *ETV4* (ETS variant transcription factor 4) and *SPRY4* (Table S3). ETV4 is a downstream target of ERK signaling pathway. In a mouse model, ETV4 promotes prostate cancer metastasis in response to coactivation of PI3-kinase and Ras signaling pathways (Aytes et al., 2013). Our results suggest that expression levels of *SPRY2* and *ETV4* are likely indicative of the sensitivity of cancer cell lines to many MAP kinase inhibitors. Other examples of drugs whose most frequently selected genes matched the drug-target genes include venetoclax-*BCL2*, navitoclax-*BCL2*, daporinad-*NAMPT* and savolitinib-*MET* (Table S3).

#### Drugs whose top-ranked genes did not match the drug targets

Our analysis also identified predictive genes that did not match the drug targets or genes for drugs with unknown mechanism of action. Here we provide two such examples.

First, *UGT1A10* (UDP glucuronosyltransferase family 1 member A10) was the gene most frequently selected as predicting sensitivity to the drugs luminespib (HSP90 inhibitor), PI-103 (mTOR inhibitor), PCI-34051 (HDAC inhibitor), and UMI-77 (MCL1 inhibitor). UGT1A10 is an enzyme of the glucuronidation pathway that is involved transforming drugs into water-soluble, excretable metabolites for drug detoxification (Dellinger et al., 2007; Dellinger et al., 2006). UGT1A10 is also thought to act as an antioxidant because of its detoxification function. However, to our best knowledge, neither HSP90, mTOR or luminespib has been linked to UGT1A10.

Second, *TRPM4* (transient receptor potential melastatin 4) was selected in 100% of gene sets that predicted sensitivity to acetalax. On the other hand, *TRPM4* was selected in fewer than 5% of the sets for all other drugs, suggesting that the *TRPM4*-acetalax interaction is specific. The mechanism of action for acetalax is unknown. Recent studies suggested that TRPM4 may be implicated in regulating cancer cell migration and in the epithelial-to-mesenchymal transition (Gao and Liao, 2019; Sagredo et al., 2019). *TRPM4* expression in the CCLE cancer cell lines was inversely correlated with the IC_50_ of acetalax in those cell lines (*ρ*_*S*_ = -0.45, *p*-value < 2.2E-16) (Figure S4), suggesting that cancer cell lines with higher *TRPM4* expression are more sensitive to acetalax.

### Predicting *In-vivo* Drug Sensitivity Based on *in-vitro* Data: Proof of Concept

If the transcriptome of a cancer cell line is predictive of that cell line’s sensitivity to a drug, we hypothesize that the transcriptome of a corresponding normal tissue is also predictive of that tissue’s sensitivity to the drug. To probe this hypothesis, we chose trametinib (GDSC drug ID=1372, a MEK1/MEK2 inhibitor) (Lugowska et al., 2015) as an example. From the cell-line data, we observed that the ranks of the observed IC_50_ values of trametinib in the 571 cancer cell lines were associated with the corresponding IC_50_ values predicted by the cell-line gene expression levels (*ρ*_*S*_ = 0.635, *p*-value < 1E-16) (Figure S1). Trametinib specifically binds to and inhibits MEK1 and MEK2, resulting in an inhibition of growth factor-mediated cell signaling and cellular proliferation in various cancers (Lugowska et al., 2015). Trametinib is clinically approved for treating all stages of melanoma and tumors with the BRAF V600E mutation including colorectal cancer (Bedard et al., 2015; Corcoran et al., 2015; Grob et al., 2015; Long et al., 2015; Robert et al., 2015). GDSC assays also demonstrated that cancer cell lines from the skin and intestine are especially sensitive to trametinib (Iorio et al., 2016) compared to those from other organs.

Using the cell-line data as the training data, we predicted the IC_50_ values of trametinib for the ∼11,000 TCGA RNA-seq samples. Our results indicated that the median predicted IC_50_ values of trametinib for melanoma (both skin and uveal) and intestine tumors (colorectal and rectal,) were much lower (showing higher sensitivity to trametinib) than those for all other tumor types (Figure 2A). Those results are consistent with both trametinib’s specificity for cancer cell lines derived from those tissues (Iorio et al., 2016) and its clinical efficacies; these consistencies suggest that our approach of linking *in-vitro* and *in-vivo* drug sensitivities is sound. Trametinib is effective in treating patients with colorectal tumors with BRAF V600E mutations (Bedard et al., 2015; Corcoran et al., 2015) and melanoma (Robert et al., 2019). Moreover, our analysis further demonstrated that the median predicted IC_50_ values of trametinib are overall much lower for colorectal tumors than for the adjacent “normal” tissues (Figure 2B). Likewise, our result indicated that trametinib is less cytotoxic to most normal organs except blood and spleen (Figure 2B), both of which are hematopoietic related. Such tumor-to-normal selectivity is not common among the 272 drugs (Table S5; see below ‘Tumor-to-normal sensitivity’). It is also reassuring that the predicted IC_50_ values of trametinib for normal intestines from GTEx are comparable to those for the TCGA “normal” colon tissues (Figure 2B). Interestingly, breast invasive carcinoma (BRCA) and prostate adenocarcinoma (PRAD) tumors are predicted to be the least sensitive to trametinib among the 33 TCGA tumor types (Figure 2A).

**Figure 2.**
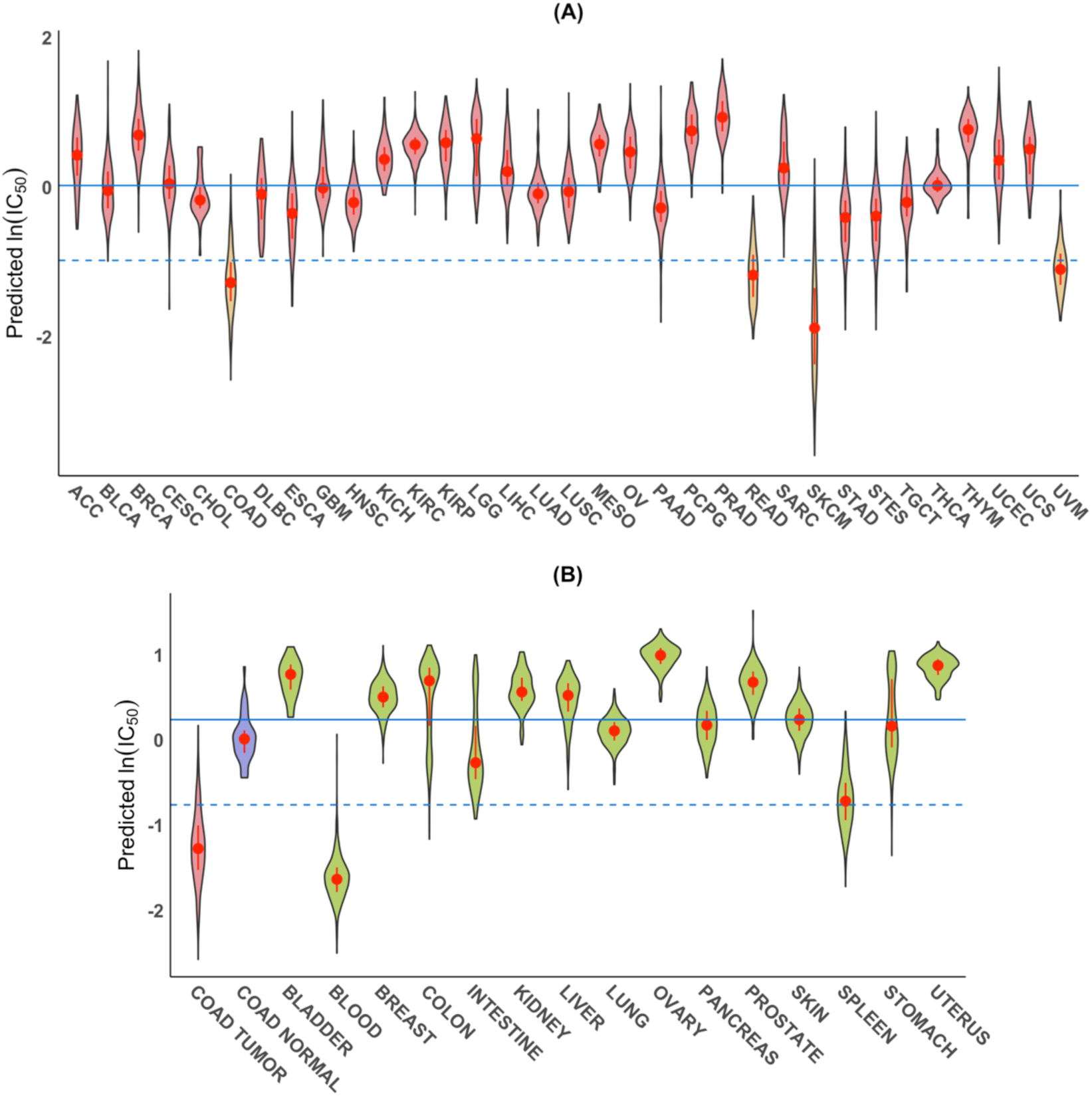
Predicted sensitivity of tumor-types and normal tissue to trametinib. (**A**) Violin plots of predicted ln(IC_50_) values of trametinib based on RNA-seq gene expression data from TCGA tumor samples from 33 tumor types. Overall COAD, READ, SKCM and UVM tumors (yellow) had the lowest predicted median IC_50_ values. For the description of the 33 TCGA tumor types, see supplementary data (Table S6). The solid line shows the median of the medians of the predicted IC_50_ values for all 33 tumor types whereas the dashed line is one logarithmic unit below the solid line. (**B**) Violin plots of the predicted ln(IC_50_) values of trametinib for COAD tumor (red) and normal (blue) samples from TCGA and for GTEx normal tissue samples from 15 major organs (green); here the solid line shows the median of the medians of the predicted IC_50_ values for all 16 normal tissues. In each violin, the red dot is located at the median; the vertical red bar extends from 25th to 75th percentiles.

Although the median predicted IC_50_ values of trametinib for samples from other tumor types were relatively high, some individual tumor samples were predicted to be as sensitive as the colorectal tumor samples to trametinib, e.g., a few of the PAAD (pancreatic adenocarcinoma) samples. The ability to predict drug sensitivity of individual tumors is important for personalized medicine.

### Predicting Sample-specific IC_50_s for All TCGA and GTEx Samples to All 272 Drugs

After establishing the potential utility of our concept, we predicted the sensitivities (IC_50_ values) of all TCGA (Table S7) and GTEx samples for the top 272 drugs using our tumor and GTEx data, respectively. Overall, most of the TCGA tumor samples were predicted to be highly sensitive (pan cancer median predicted ln(IC_50_) < 0) to about 35 of the 272 drugs (Table S5). Many of the drugs target DNA/protein synthesis, cell cycle, microtubules, and the mTOR pathway. Most of the drugs were also predicted to be similarly cytotoxic to normal samples from TCGA (Table S5). Those drugs are among the most commonly used chemotherapeutic agents. Unfortunately, they are also associated with high cytotoxicity to normal organs.

### Tumor-to-normal Sensitivity

For each of the 272 drugs, we compared the median predicted IC_50_ of the drug for all tumor samples with the median predicted IC_50_ value for all normal samples from the same tumor type from TCGA. We only considered the 14 tumor types (BRCA, COAD, HNSC, KICH, KIRC, KIRP, LIHC, LUAD, LUSC, PRAD, STAD, STES, THCA, and UCEC) with more than 20 normal samples.

We identified eight drugs whose median predicted ln(IC_50_) value for tumor samples was more than one logarithmic unit lower than that for corresponding normal samples in at least one of the 14 tumor types (Figure S5). One logarithmic unit corresponds to tumor tissue being about 2.7-fold more sensitive than normal tissues. Though we selected drugs based on at least one tumor type having this high ratio of tumor-to-normal sensitivity, for most drugs the high ratio was not limited to a single tumor type. Among the eight drugs, trametinib is an exceptional example for which a drug is predicted to not only be specific for a tumor type (COAD, in this case) but also have high tumor-to-normal sensitivity for only a single tissue type among the 14 tumor-normal pairs (Figure 3A). Similarly, luminespib (Hsp90 inhibitor) (Figure 3B) and sapitinib (Erbb inhibitor) (Figure 3C) are predicted to have high tumor specificity largely for LUSC with high tumor-to-normal sensitivity for the tissue type.

**Figure 3.**
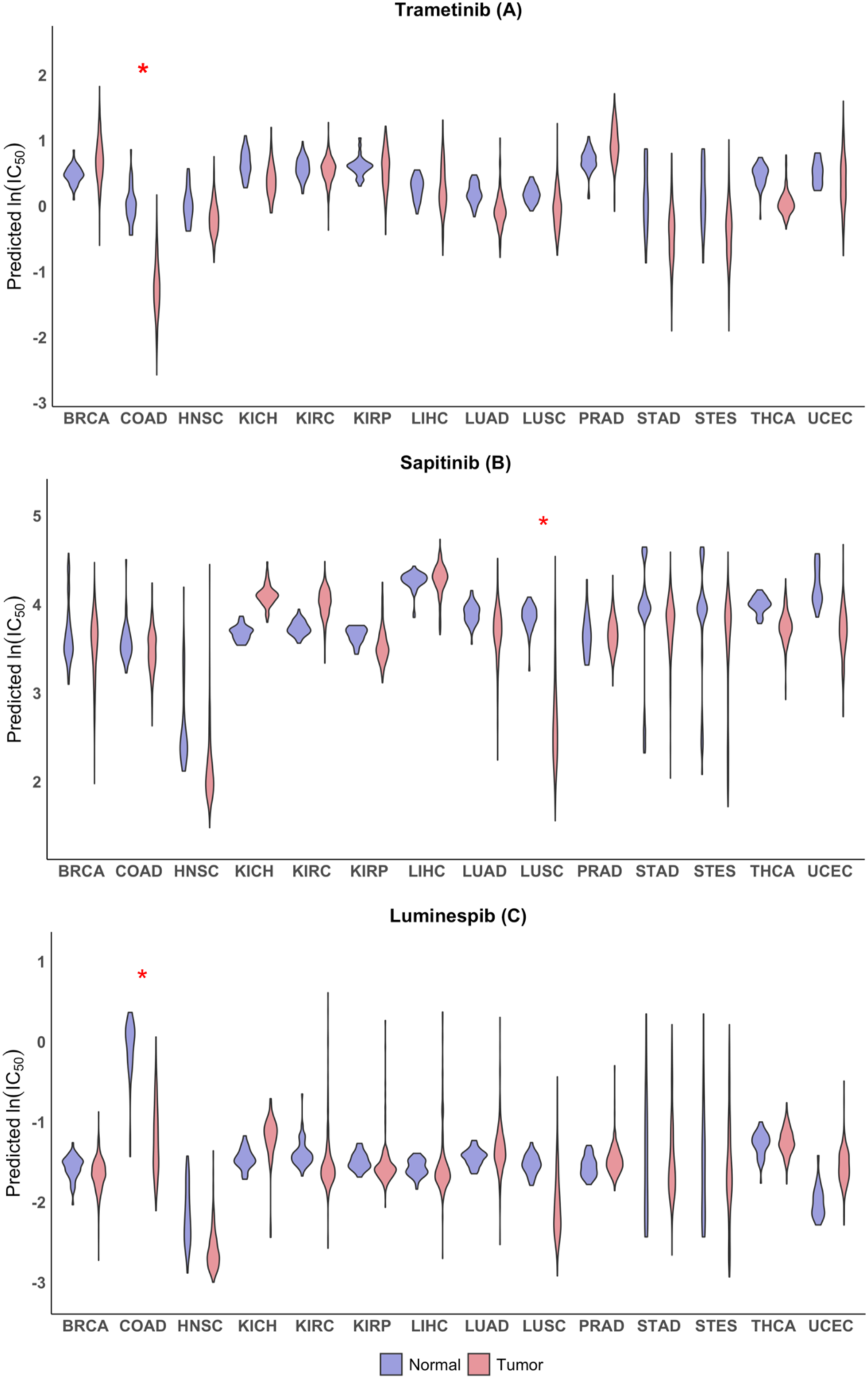
Examples of drugs that are predicted to have high tumor-to-normal sensitivity for some tumor types. Violin plots of predicted IC_50_ values in tumor (red) and normal (blue) tissue for trametinib (**A**), sapitinib (**B**) and luminespib (**C**) that showed the ratio of tumor-to-normal sensitivity exceeding 2.7 (1 logarithmic unit) for at least one of 14 tissue types. The ln(IC_50_) values of the drugs were predicted based on the RNA-seq data of the tumor and normal tissue samples from TCGA. Violin plots for normal and tumor samples from the same tissue type are shown as side-by-side pairs with their TCGA type on the X-axis. See Figure 2 legend for additional description of the violin plots. Red star (*) indicates the difference between the median of predicted IC_50_ values for normal samples and the median of predicted IC_50_ values for tumor samples is more than one logarithmic unit.

### Tumor-type-specific Drugs

For each drug, we compared its median predicted IC_50_ value among samples from one tumor type with the median of the medians of the predicted IC_50_ values from all 33 tumor types. We considered a drug to be specific for a tumor type if the median predicted IC_50_ value for the tumor type is one logarithmic unit (∼2.7 times) lower than the median of the medians from all tumor types. We identified 109 such drugs (Table S8), most (96) of which were predicted to have lower IC_50_s for either diffuse large B cell lymphoma (DLBC), thymoma (THYM) or both. Interestingly, 12 of the remaining 13 drugs that were predicted not specific for DLBC, THYM or both are kinase inhibitors, consistent with the notion that kinase inhibitors target specific cellular pathways. Eighty-three drugs were predicted to have higher specificity for a unique tumor type; 74 for DLBC, 5 for SKCM (skin cutaneous melanoma), 2 for THYM, and 1 for HNSC (head-neck squamous cell carcinoma) and 1 for KIRP (kidney renal papillary cell carcinoma). Our analysis suggested that the B-Raf proto-oncogene (BRAF) inhibitors (AZ628, Dabrafenib, PLX-4720, and SB590885) are specific for melanoma (Table S8). We also predicted that the four mitogen-activated protein kinase kinase (MEK/ERK) inhibitors (PD0325901, SCH772984, selumetinib, and trametinib) are specific for both colorectal cancer and melanoma. Indeed, clinical trials have demonstrated clinical efficacy of BRAF inhibitors for a portion of melanoma patients harboring activating BRAF mutations (Alcala and Flaherty, 2012; Corcoran et al., 2015; Long et al., 2015; Robert et al., 2019; Robert et al., 2015). Thus, our predictions are consistent with those human clinical trial results.

We predicted that acetalax, with unknown mechanism of action, was specific for multiple tumor types including prostate adenocarcinoma (PRAD) and breast invasive carcinoma (BRCA) (Figure 4A, Table S8). We predicted alisertib was specific for DLBC and lower-grade glioma (LGG) (Figure 4B). Several tumor types including MESO (mesothelioma) and OV (ovarian serous cystadenocarcinoma) were predicted to be highly sensitive to dasatinib (Figure 4C). We predicted dabrafenib to be specific for DLBC and SKCM (Figure 4D). OSI-027 (mTOR inhibitor) showed high specificity to BRCA and PRAD (Figure 4E). Sapitinib (EGFR/HER2 inhibitor) was specific for HNSC and cervical squamous cell carcinoma and endocervical adenocarcinoma (CESC), esophageal carcinoma (ESCA), and lung squamous cell carcinoma (LUSC) (Figure 4F).

**Figure 4.**
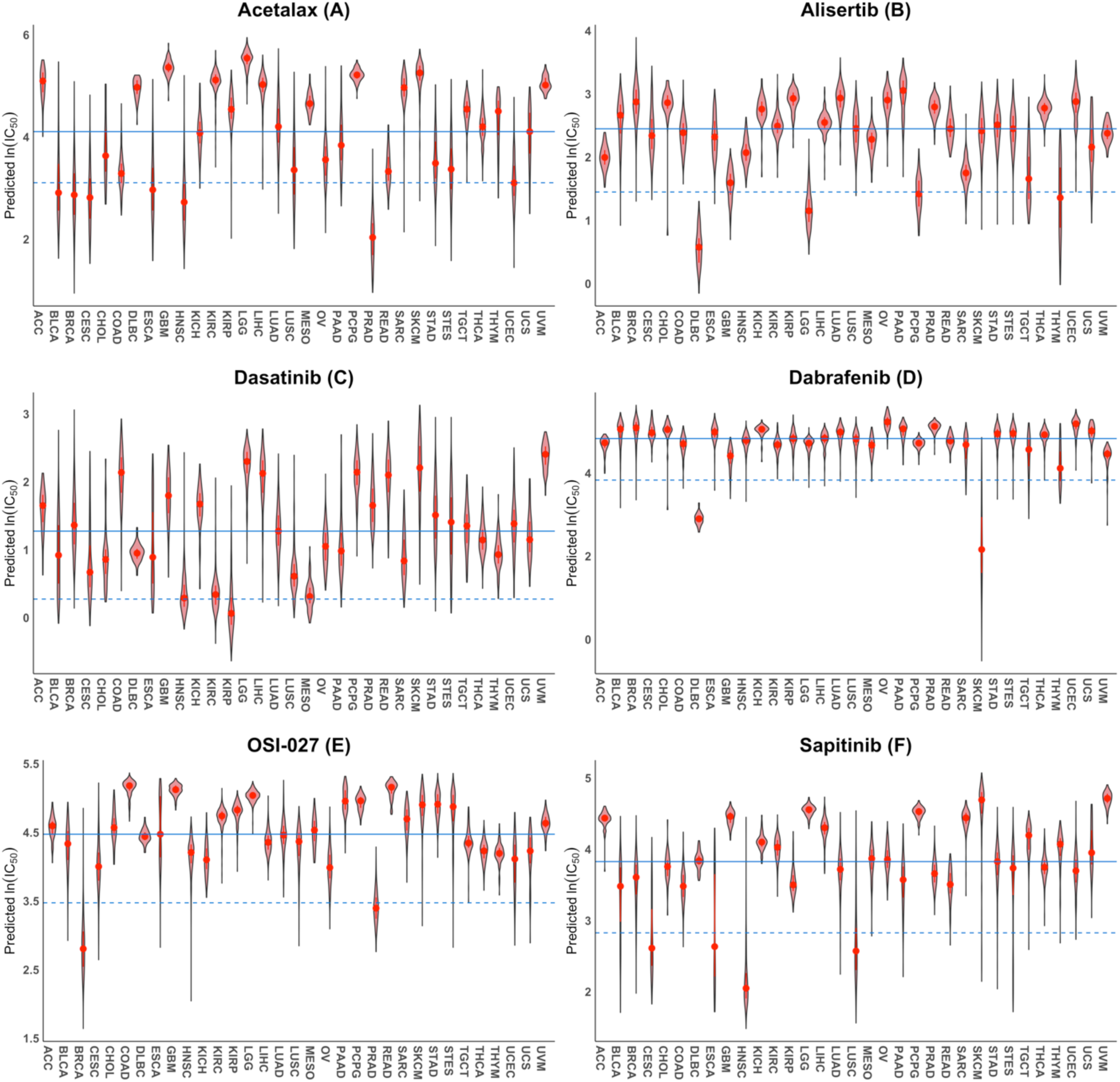
Selected drugs that are predicted to be tumor-type-specific. Violin plots of the predicted ln(IC_50_) values of Acetalax (**A**), Alisertib (**B**), Dasatinib (**C**), Debrafenib (**D**), OSI-027 (**E**), and Sapitinib (**F**) for TCGA tumor samples from 33 tumor types. The solid line shows the median of the medians of the predicted IC_50_ values for all 33 tumor types; whereas the dashed line is one logarithmic unit below the solid line. See Figure 2 legend for additional description of the violin plots.

### Drug Sensitivity of Breast Cancer Subtypes

Breast cancers may be classified into subtypes bases gene-expression signatures (Sotiriou and Pusztai, 2009). To see if subtypes of breast cancer were predicted to show differential sensitivity to any of the 270 drugs, we divided the ∼1,100 TCGA BRCA samples into five subgroups (basal-like, Her2-positive, luminal A, luminal B, and normal-like) based on the PAM50 classification (Chia et al., 2012; Wallden et al., 2015). For each subtype, we compared the median of the predicted IC_50_ values of a drug for the samples of the subtype with the median of the medians of the predicted IC_50_ values for the five subtypes. We focused on drugs for which the difference in the medians exceeded 0.5 logarithmic units (corresponding to a 1.7-fold difference in IC_50_). Among the 270 drugs, seven drugs met this criterion (Table 1). Although OSI-027 did not meet the criterion, we also included it in Table 1 as it is the only drug among the top 272 drugs that showed the highest overall specificity for breast cancer compared to all other TCGA tumor types (Figure 4E). We showed that Her2-positive breast cancer subtype is predicted to have higher sensitivity to OSI-027 compared to all other four subtypes. We also predicted that basal-like subtype breast cancer has higher sensitivity to five (bleomycin, daporinad, sepantronium bromide, etoposide, and ICL1100013) of the seven drugs, luminal B subtype breast cancer has higher sensitivity to ABT737 and navitoclax, both of which are BCL2 inhibitors.

Bleomycin is effective for elderly patients with metastatic breast cancer (Campana et al., 2014). Bleomycin sulfate followed by electroporation treatment in patients with recurrent in-breast or chest-wall tumors is effective (Paramanov et al., 2007). We predicted that bleomycin has the highest sensitivity for basal-like breast cancer among the three subtypes (Figure 5A). Interestingly, the most frequently selected gene for predicting sensitivity to bleomycin was *ACE* (Table S3). The *ACE* gene encodes the angiotensin I converting enzyme. Although the exact mechanism of action for bleomycin is unclear, it is thought to inhibit DNA synthesis. *ACE* expression in cancer cell lines was positively correlated with the IC_50_ value of bleomycin in those cell lines (Figure 5B), suggesting that higher *ACE* expression in cancer cell lines is associated with higher resistance to bleomycin. Although the literature on the relationship between bleomycin and ACE is limited, Day et al. (Day et al., 2001) demonstrated that treatment of primary bovine pulmonary artery endothelial cells with bleomycin did increase ACE enzymatic activity and *ACE* mRNA and that the increased ACE expression resulted in fibrosis. Mechanistically, bleomycin activated p42/p44 MAP kinase which in turn up-regulated EGR1, a transcription factor that positively regulates *ACE* expression (Day et al., 2001). Bleomycin-induced ACE overexpression can be inhibited using MEK1/2 inhibitors (Day et al., 2001). Similarly, Li et al. reported that inactivation of ACE alleviated bleomycin-Induced lung Injury (Li et al., 2010). Those studies clearly establish a link between bleomycin and ACE in fibrosis. Interestingly, patients treated with ACE inhibitors have a lower than expected chance of developing cancer (Rosenthal and Gavras, 2009). Both fibrosis (Chandler et al., 2019) and MAP kinase activation (Dhillon et al., 2007) are associated with tumor progression. Taken together, those results suggest that combination therapy using bleomycin and MEK and/or ACE inhibitors could be beneficial for treating cancers, particularly basal-like type breast cancer.

**Figure 5.**
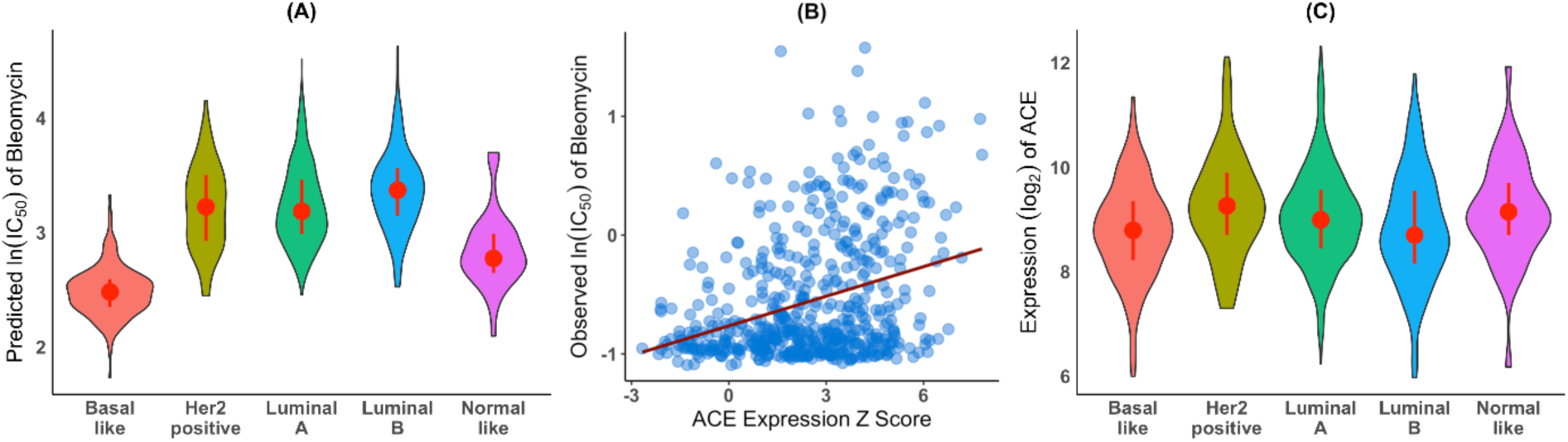
Basal breast tumors are predicted to be more sensitive to bleomycin than luminal A, luminal B or Her2-positive breast tumors and the sensitivity is inversely correlated with *ACE* expression. (**A**) Predicted bleomycin sensitivity for the five subtypes of TCGA BRCA samples: violin plots of the predicted ln(IC_50_) values of bleomycin for the five subtypes of breast tumors based on gene expression data and PAM50 classification of TCGA BRCA samples. (**B**) *ACE* gene expression in cancer cell lines versus sensitivity to bleomycin: *ACE* expression in the CCLE cancer cell lines was positively correlated with observed ln(IC_50_) values for bleomycin (*ρ*_*S*_ = 0.27, p-value = 6.6E-11). The red line is the least-squares regression line. (**C**) TCGA breast cancer tumor gene expression data: violin plots of *ACE* expression in TCGA basal-like, Her2-positive, luminal A, luminal B, and normal-like breast tumor samples.

*ACE* expression levels in the five breast cancer subtypes in TCGA tumor samples were similar (Figure 5C). This similarity may not be surprising as the TCGA tumor samples were taken before any chemotherapy, thus, no induction of *ACE* expression by bleomycin. Our results together with the relationship between bleomycin and *ACE* expression described in the preceding paragraph suggest that patients with basal-like breast tumors would be more sensitive to bleomycin; and if treated with bleomycin, would have lower *ACE* expression, thus, less fibrosis, than similarly treated patients with the luminal subtypes and Her2-positive subtype.

We further found that among the five subtypes of breast cancer Her2-positive subtype had the highest sensitivity to OSI-027 (Figure 6A). OSI-027 is a potent inhibitor of mTOR1 and mTOR2 (Bhagwat et al., 2011). In a preclinical trial, OSI-027 showed significant inhibition of tumor growth in several human cancer xenograft models (Bhagwat et al., 2011).

**Figure 6.**
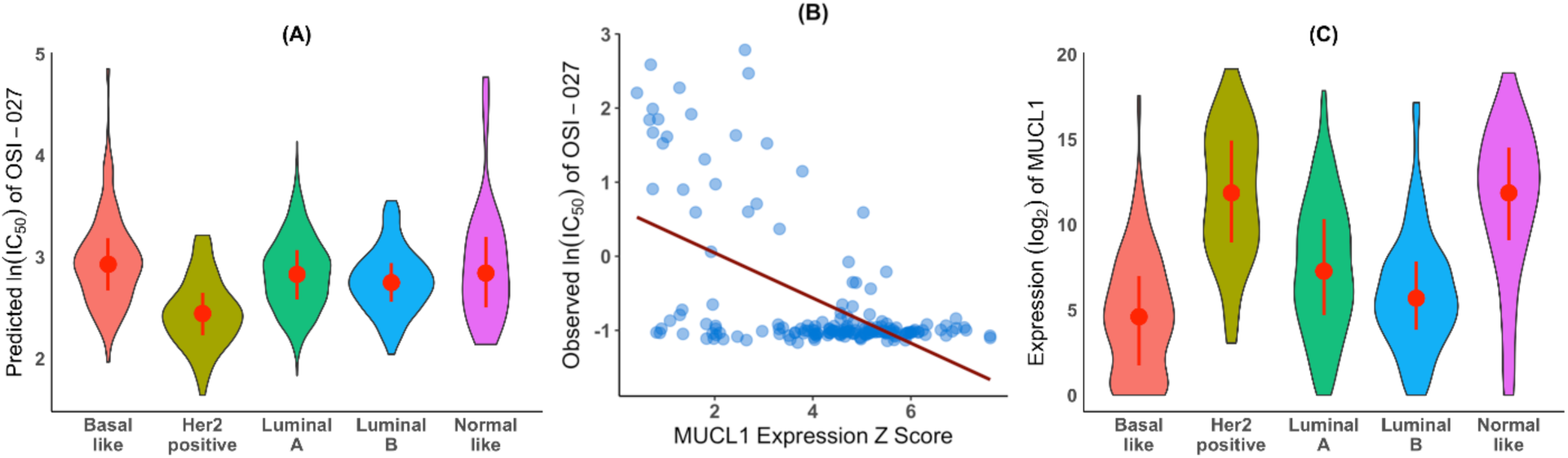
Breast cancer Her2-positive subtype was predicted to have the highest sensitivity to OSI-027 (GDSC drug ID=1594) and that this sensitivity may be directly linked with *MUCL1* expression level in those tumors. (**A**) Predicted OSI-027 sensitivity for the five subtypes of TCGA BRCA samples: violin plots of the predicted ln(IC_50_) values of OSI-027 for basal-like, Her2-positive, luminal A, luminal B and normal-like breast cancer tumors. (**B**) *MUCL1* gene expression in cancer cell lines versus sensitivity to OSI-027: *MUCL1* expression in CCLE cancer cell lines was negatively correlated with the observed ln(IC_50_) of OSI-027 (*ρ*_*S*_ = –0.30, *p* = 6.0E-05), indicating higher *MUCL1* expression was associated with higher sensitivity to OSI-027. (**C**) TCGA breast cancer tumor gene expression data: violin plots of *MUCL1* expression in TCGA basal-like, Her2-positive, luminal A, luminal B, and normal-like breast tumor samples.

*MUCL1* (mucin like 1) was the most frequently selected gene for predicting sensitivity to OSI-027 (Table S3). *MUCL1* expression in CCLE cancer cell lines was inversely correlated with the observed IC_50_ of OSI-027 for those cancer cell lines (*ρ*_2_=-0.30, *p*-value=6.0E05) (Figure 6B), suggesting that cancer cell lines with higher *MUCL1* expression had higher sensitivity to OSI-027. *MUCL1* expression in TCGA Her2-positive breast tumors was also significantly higher than in any other four subtype breast tumors (*p*-value=5.42e-09, Mann-Whitney-Wilcoxon) (Figure 6C), suggesting that *MULC1* expression level may be a surrogate biomarker of the effectiveness of OSI-027 for breast tumors. *MUCL1* is a breast-specific gene expressed in normal and tumor human mammary epithelium (Colpitts et al., 2002; Miksicek et al., 2002). *MUCL1* is regulated by Her2 and is implicated in breast cell proliferation through the FAK/JNK pathway (Conley et al., 2016). *MUCL1* expression level in breast cancer is associated with Her2 over-expression (Valladares-Ayerbes et al., 2009). Our predictions indicate that Her2-positive subtype of breast tumors was most sensitive to OSI-027 and that this sensitivity may be directly linked with *MUCL1* expression level in those tumors.

## DISCUSSION

In this work, we began by investigating if a cancer cell line’s transcriptome (Barretina et al., 2012; Ghandi et al., 2019) can predict the IC_50_ of a drug acting on that cell line for each of the 473 GDSC drugs and 1,019 cell lines (Garnett et al., 2012; Iorio et al., 2016; Yang et al., 2013). We found that, for about half of the drugs, transcriptomes were reasonably predictive of the sensitivity of the cell lines to those drugs, *i*.*e*., that Spearman correlation between predicted and observed IC_50_ values >0.4.

Among those drugs for which gene expression data can reasonably predict a cancer cell line’s sensitivity, we identified many known drug-gene interactions as well as several novel associations. Our results are consistent with and lend additional support to the notion that the expression levels of *ABCB1* and *SLFN11* are potential biomarkers for cancer cell line sensitivity to multiple drugs (Ballestrero et al., 2017; Chen and Sikic, 2012; Pietanza et al., 2018; Vaidyanathan et al., 2016). Our results also revealed that *SPRY2* expression is positively correlated with the sensitivity of the cancer cell lines to many MEK inhibitors from GDSC, suggesting that *SPRY2* expression may be a predictive biomarker for the effectiveness of MEK kinase inhibitors. We also uncovered that *C19orf33* expression in cancer cell lines is positively correlated (*ρ*_2_ ≥ 0.3) with the IC_50_ values of hundreds of chemotherapeutic drugs in those cell lines (Table S4), suggesting that *C19orf33* expression may be a general biomarker for cancer cell line sensitivity to chemotherapeutic agents.

We applied the predictive models learned from the cell line data to both tumor and normal TCGA and normal GTEx RNA-seq data. We used trametinib, a MEK kinase inhibitor (Lugowska et al., 2015), as a proof-of-concept exemplar. Based on the GDSC assay data, cancer cell lines from the skin and intestine had the highest sensitivity to trametinib. Clinically, trametinib has been approved for treating cancer patients with the BRAF V600E mutation (Bedard et al., 2015; Corcoran et al., 2015; Grob et al., 2015; Long et al., 2015; Robert et al., 2015). Our analyses of the TCGA tumors revealed that trametinib has the highest specificity for melanoma, colorectal cancer, and rectal cancer among all 33 TCGA tumor types, consistent with the clinical application of trametinib. This result prompted us to extend our predictions to the top 272 predictable GDSC drugs, those for which a cancer cell line’s sensitivity can be predicted from transcriptome data.

We found that some of the drugs are highly cytotoxic to all tumors from all tumor types. Those drugs were also predicted to be cytotoxic to normal organ tissues. Unfortunately, those drugs are among the most frequently used chemotherapeutic agents and are associated with side effects. We also identified drugs that show higher specificity towards one or a few tumor types, e.g., ERK, MEK and BRAF inhibitors (AZ628, dabrafenib, PD0325901, PLX-4720, SB590885, SCH772984, and trametinib) for melanoma and colorectal cancer, ERBB/EGFR inhibitors (afatinib and sapitinib) for head-neck squamous cell carcinoma, and BCL2 inhibitors (ABT737, navitoclax and venetoclax) for low grade glioma and glioblastoma multiforme. We further uncovered that OSI-027 was highly specific for breast cancer, especially the Her2-positive subtype breast cancer tumors. Furthermore, our result suggests that *MUCL1* expression may be a surrogate marker for tumor response to OSI-027.

We also predicted that a few drugs not only were tumor-type specific but also induced higher sensitivity in tumor tissue than normal tissue for those tumor types. Those drugs may have better clinical efficacies. Our result suggested that paclitaxel (microtubule inhibitor) (commonly known as taxol) was more specific for breast, lung, and uterine tumors than other tumor types and that those tumors were overall more sensitive to paclitaxel than corresponding normal tissue. Similarly, we predicted that trametinib was specific for melanoma, colorectal tumors and rectal tumors; our analysis also found that those tumors were also more sensitive to trametinib than the normal tissues of the same origins. Sapitinib (ERBB inhibitor) was predicted to have the highest specificity for lung squamous cell carcinoma and was also predicted to be more cytotoxic to lung carcinoma than to normal lung tissue.

We used breast cancer as example to identify tumor subtypes that may be especially sensitive to a drug. For example, we predicted that five drugs (bleomycin, daporinad, sepantronium bromide, etoposide, and ICL1100013) are more specific for basal-like breast cancer subtypes whereas ABT737 and navitoclax sapitinib and afatinib are more specific for the luminal subtypes especially luminal B. We predicted that OSI-027 (mTOR inhibitor) is specific for breast cancer among all 33 TCGA tumor types, especially the Her2-positive breast cancer subtype. Since we have provided the predicted IC_50_ values for all TCGA tumor samples (Table S7), our approach can be easily applied to other tumor types.

It is worth pointing out that OSI-027 was assayed twice (GDSC1 by the Massachusetts General Hospital and GDSC2 by the Sanger) with two different drug IDs (299 for GDSC1 and 1594 for GDSC2). GDSC1 screened 906 cancer cell lines for the drug; whereas GDSC2 screened 265 cancer cell lines. The IC_50_ values of OSI-027 for breast cancer cell lines from GDSC2 are the lowest among all 265 cancer cell lines; however, OSI-027 IC_50_s for breast cancer cell lines from GDSC1 were not among the lowest. Consequently, we predicted that OSI-027 with drug ID 1594, but not drug ID 299, is specific for breast cancer. Since GDSC recommends using assay data from GDSC2 when available, our results for OSI-027 may still be valid. However, additional replications are warranted.

There are many challenges associated with relating findings from cell lines to tumors and clinical applications. For example, *in vitro* assays do not capture organ responses. Although we used the largest collection of cancer cell lines in our predictive models, our models have limitations when applied to datasets that may not be represented by the training data. Nonetheless, translating findings from cell lines to tumors has had some success (Barretina et al., 2012; Geeleher et al., 2014; Geeleher et al., 2017; Rajapakse et al., 2018; Reinhold et al., 2012; Reinhold et al., 2019).

Our method uses the *k*-nearest neighbor rule to predict drug response of an unknown sample. The predicted value of a sample is taken as the average of the values of its *k*-nearest neighbors. Because of the averaging, the most extreme predicted values, either high or low, usually cannot be as extreme as the corresponding observed values. Therefore, although the correlation between the predicted and observed values can be high, e.g., 0.8, the magnitude of the predicted values is generally pulled in from the extremes; the trend of the predicted values among the samples, however, is usually preserved (Figure S1). This information should be kept in mind when interpreting the “face value” of predicted values.

In summary, our predicted drug sensitivity data for all TCGA tumor and normal samples should be a valuable resource to researchers and clinicians. We identified known and novel drug-gene interactions and potential biomarkers for drug effectiveness. Our approach is unique in that we not only predicted drug specificity for tumor types and subtypes, but also drug sensitivity towards normal tissues. We predicted a few drugs to have high specificity for some tumor types compared to all others and high ratios of tumor-to-normal sensitivity. If true, our predictions could have clinical relevance for patients’ care.

## ACKNOWLEDGEMENTS

We are grateful to the NIEHS Office of Scientific Computing for computing support and also to Dr. Frank Day for help with the website. The PAM50 classification of the TCGA breast cancer tumor samples was kindly provided by Joel Parker (UNC). This research was supported by the Intramural Research Program of the National Institutes of Health, National Institute of Environmental Health Sciences (ES101765).

## AUTHOR CONTRIBUTIONS

Conceptualization and methodology, L.L.; Research and investigation: Y.L., L.L., D.M.U., I.S., X.L.; Algorithm optimization, J.M.K.; Writing, L.L., Y.L., D.M.U. with help from all authors.

## EXPERIMENTAL PROCEDURES

### Data

#### Drug sensitivity of cancer cell lines

GDSC screened 397 distinct compounds and 1,000 distinct cell lines in two releases: GDSC1 screened 320 compounds in 987 cell lines whereas GDSC2 screened 175 compounds in 809 cell lines. GDSC reports the ln(IC_50_) for each combination of cell line and compound as a measure of the sensitivity of cell viability in that cell line to the compound. We downloaded the ln(IC_50_) data from the GDSC website https://www.cancerrxgene.org/downloads/bulk_download. When combining data from both releases, if IC_50_s for the same cell line and compound were present in both, we kept only the one from GDSC2 as advised by GDSC. In total, 453 drugs were assayed, among which 397 were unique (56 had two different drug IDs). For those with two unique drug IDs, we did not combine them but rather treated each as it were a unique drug as GDSC did on their website. Like GDSC, we refer to the ln(IC_50_) values simply as IC_50_ throughout the manuscript, unless specified otherwise.

#### Gene expression of cancer cell lines

CCLE measured gene expression profiles using RNA-seq for 1,019 cancer cell lines. We downloaded the gene expression data from the CCLE website (https://portals.broadinstitute.org/ccle/) (CCLE_RNAseq_rsem_genes_tpm_20180929.txt) and converted Ensembl gene IDs into official gene symbols using the annotation file (gencode.v19.genes.v7_model.patched_contigs.gtf). For 111 genes (primarily small nucleolar genes), multiple Ensembl entries corresponded to the same gene symbol; we used the average expression value for those genes.

#### Gene expression of tumor tissue

TCGA makes available RNA-seq gene expression profiles from 771 normal and 10,339 tumor samples encompassing 33 tumor types (Table S6). We downloaded the RSEM-normalized expression data from the Broad GDAC firehose (https://gdac.broadinstitute.org/). We then log2-transformed those expression values after adding 1 to each.

#### Gene expression of normal tissue

GTEx has available RNA-seq gene expression data for normal tissue samples. We downloaded these data (GTEx_Analysis_2017-06-05_v8_RNASeQCv1.1.9_gene_tpm.gct.gz) from the GTEx website https://www.gtexportal.org/home/datasets ; we extracted RNA-seq expression data for 5,894 tissue samples from 15 major organs (bladder, blood, breast, colon, intestine, kidney, liver, lung, ovary, pancreas, prostate, skin, spleen, stomach, and uterus). We similarly transformed the data as above.

#### Combining GDSC drug sensitivity and CCLE gene expression for cancer cell lines

Among the cell lines used by GDSC, we identified all those for which CCLE provided gene expression profiles. Accordingly, for each of the 453 drugs from GDSC, we have CCLE gene expression profiles for a subset of the cell lines with IC50s for that drug. Denote the number of such cell lines for drug *D* by *N*_*D,CCLE*_ and the number of genes in the expression profile by *G* (same for every drug, *G* = 19163). We created a gene expression data matrix (*G* × *N*_*D,CCLE*_) for each drug, with each row indexing a gene and each column indexing a cell line. We also created a corresponding drug-specific vector of IC_50_ values (with length *N*_*D,CCLE*_). Here *N*_*D,CCLE*_ ranged from 38 to 579 with 25^th^, 50^th^, and 75^th^ percentiles of 473, 538, and 553, respectively. For clarity, we refer to these data matrices as “cell-line data”.

#### Combining GDSC drug sensitivity and CCLE gene expression with TCGA or GTEx gene expression

We augmented each of the 453 matrices of cell-line data with columns of RNA-seq expression profiles for the tumor samples from the TCGA using the common genes between the two (*G* = 19,163). Thus, we created 400 new expression data matrices (*G* × *N*_*D,CCLE*+*TCGA*_), one for each drug. Here *N*_*D,CCLE*+*TCGA*_ ranged from 11,129 to 11,670 with 25^th^, 50^th^, and 75^th^ percentiles of 11,563, 11,629, and 11,644, respectively. We refer to these data matrices as “tumor data”.

Similarly, we augmented each of the 453 matrices of cell-line data with columns of RNA-seq expression profiles for the normal tissue samples from GTEx using the common genes between the two (*G* = 19163). We created an additional 400 expression data matrices (*G* × *N*_*D,CCLE*+*GTEx*_), one for each drug. Here *N*_*D,CCLE*+*GTEx*_ ranged from 5,932 to 6,473 with 25^th^, 50^th^, and 75^th^ percentiles of 6,367, 6,970, and 6,447, respectively. We refer to these data matrices as “GTEx data”.

#### TCGA breast tumor sample clinical data

Hormone status of the TCGA breast invasive carcinoma (BRCA) tumor samples (file name: BRCA.clin.merged.txt) was downloaded from the Broad GDAC firehose (https://gdac.broadinstitute.org/).

### Data Integration

When combining data from different sources, it is important that the data are comparable. For this purpose, we computed Z-scores across genes for each cell line from CCLE and each sample from TCGA or GTEx (referred to collectively as “samples”). Thus, each sample has a mean expression of 0 and standard deviation of 1. Let *x*_*i,j*_ and *z*_*i,j*_ be the log_2_-transformed expression values before and after Z-transformation, respectively, for *j*^th^gene in *i*^th^ sample, that is,

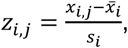

where *i* = 1, ⋯ *N*_*D,CCLE*+*TCGA*_ *or N*_*D,CCLE*+*GTEx*_ (depending on the data set) and *j* = *i*, ⋯, *G* and 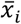 and *s*_*i*_ are the mean and standard deviation of the expression values for sample *i*.

## METHODS

### The GA/KNN Algorithm

The GA/KNN (genetic algorithm/*k*-nearest neighbors) algorithm combines a genetic algorithm for feature selection and the *k*-nearest neighbor method for classification or prediction (Li et al., 2001). In the present context, the main idea of the GA/KNN algorithm is to use an evolutionary algorithm to select many sets of *d* genes (see below) whose expression levels can accurately predict observed IC_50_ values using the *k*-nearest-neighbors prediction rule. The prediction rule is simple: the predicted IC_50_ value of a sample is defined as the average of the observed IC_50_ values of its *k* nearest neighbor samples (excluding itself) as determined by Euclidean distance in the *d*-dimensional space defined by a gene set. In making predictions for a testing-set sample (see below), we considered only samples within the corresponding training set as potential neighbors.

The GA/KNN algorithm is designed to optimize, either minimize or maximize, an objective function. For prediction, a typical objective function being minimized is the sum of the squared deviations between the observed and predicted IC_50_ values across all training samples (i.e., squared-error loss). The squared-error 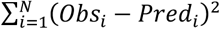, where *N* is the number of samples in the training set. The GA/KNN algorithm applied here sought a set of *d* genes to minimize this loss function.

The main parameters for the GA/KNN algorithm were set to be the same for the analyses of all datasets (Table S9). To identify the optimal number of nearest neighbors (*k*) and the “chromosome” length *d*, we systematically evaluated 16 combinations of *k* (*k*=1, 3, 5, and 7) and *d* (*d*=10, 20, 30, and 40) (Table S10). Consistent with our earlier findings (Li et al., 2001), *k*=3 and *d*=30 seemed to provide the near-optimal performance and is computationally efficient.

Because GA/KNN is computationally intensive, we only carried out 100 independent runs for each drug. To see if 100 runs were sufficient, we also analyzed trametinib with 1,000 runs. The results from both runs were comparable (Table S11 and Figure S6).

### Training and Cross-validation

For high dimensional data, multiple sets of *d* genes that can deliver similar near-optimal performance. To identify multiple sets of predictive genes, the GA/KNN algorithm uses a Monte Carlo cross-validation procedure (Xu and Liang, 2001). For each drug separately, we randomly partitioned its cell-line data into a training set (90%) and a testing set (10%). We used the training data to identify a set of *d* (*d*=30) genes whose expression levels were best predictive of the IC_50_ values of samples in the training set using a leave-one-out cross-validation procedure (Li et al., 2001). That set of *d* genes was subsequently used to predict the IC_50_ values of the testing-set samples. The average IC_50_ value of the *k*-nearest (*k*=3) training neighbors of a testing sample was taken as the predicted IC_50_ value for the testing sample. The above procedure was repeated 100 times independently, each started with a new random partition into training and testing sets. Over the 100 random training-testing partitions for a given drug, each sample would be expected to appear in about 90 training sets and about 10 testing sets. The final predicted value for a training-set sample was the average of the predicted IC_50_ values for that sample over the subset of the 100 independent training-testing partitions in which that sample appeared in a training set; analogously, the final predicted value for a testing-set sample was the average predicted IC_50_ value over the partitions where that sample appeared in a testing set.

### Assessing the Importance of Individual Gene’s Expression Levels to Prediction

Because each training-testing partition provided a set of 30 genes as predictors, we used the frequency with which a gene was selected into the 100 sets of 30 predictor genes as a measure of the importance of that gene in prediction.

### Identifying Predictable Drugs

We computed both the Pearson (*ρ*_*P*_) and Spearman (*ρ*_*S*_) correlation coefficients between the observed and predicted IC_50_ values for samples in the training and testing sets, respectively. We designated those drugs whose

*ρ*_*P*_ and *ρ*_*S*_ values were both greater than or equal to 0.4 (*ρ*_*P*_ ≥ 0.4 and *ρ*_2_ ≥ 0.4) for the testing set samples as predictable drugs.

### Predicting IC_50_ Values of TCGA Tumor Samples and GTEx Normal Tissue Samples

In these analyses, we only considered the 272 predictable drugs identified from the cell-line data. For each of the 272 drugs, we repeated the same GA/KNN procedure applied to the cell-line data to both the tumor and the GTEx data. Specifically, for each of the 272 drugs, we randomly partitioned the part of the tumor data from the CCLE cell lines into a training set (90%) and a testing set (10%), repeating the partitioning 100 times, as above. In addition, with each partition, we treated all TCGA samples in the tumor data and the GTEx samples in the GTEx data as additional “testing” samples for prediction.

## REFERENCES

Alcala, A.M., and Flaherty, K.T. (2012). BRAF inhibitors for the treatment of metastatic melanoma: clinical trials and mechanisms of resistance. Clin Cancer Res 18, 33–39.

Anderson, M.A., Deng, J., Seymour, J.F., Tam, C., Kim, S.Y., Fein, J., Yu, L., Brown, J.R., Westerman, D., Si, E.G., et al. (2016). The BCL2 selective inhibitor venetoclax induces rapid onset apoptosis of CLL cells in patients via a TP53-independent mechanism. Blood 127, 3215–3224.

Aytes, A., Mitrofanova, A., Kinkade, C.W., Lefebvre, C., Lei, M., Phelan, V., LeKaye, H.C., Koutcher, J.A., Cardiff, R.D., Califano, A., et al. (2013). ETV4 promotes metastasis in response to activation of PI3-kinase and Ras signaling in a mouse model of advanced prostate cancer. Proc Natl Acad Sci U S A 110, E3506–3515.

Azuaje, F. (2017). Computational models for predicting drug responses in cancer research. Brief Bioinform 18, 820–829.

Azuaje, F., Kaoma, T., Jeanty, C., Nazarov, P.V., Muller, A., Kim, S.Y., Dittmar, G., Golebiewska, A., and Niclou, S.P. (2018). Hub genes in a pan-cancer co-expression network show potential for predicting drug responses. F1000Res 7, 1906.

Ballestrero, A., Bedognetti, D., Ferraioli, D., Franceschelli, P., Labidi-Galy, S.I., Leo, E., Murai, J., Pommier, Y., Tsantoulis, P., Vellone, V.G., et al. (2017). Report on the first SLFN11 monothematic workshop: from function to role as a biomarker in cancer. J Transl Med 15, 199.

Barretina, J., Caponigro, G., Stransky, N., Venkatesan, K., Margolin, A.A., Kim, S., Wilson, C.J., Lehar, J., Kryukov, G.V., Sonkin, D., et al. (2012). The Cancer Cell Line Encyclopedia enables predictive modelling of anticancer drug sensitivity. Nature 483, 603–607.

Bedard, P.L., Tabernero, J., Janku, F., Wainberg, Z.A., Paz-Ares, L., Vansteenkiste, J., Van Cutsem, E., Perez-Garcia, J., Stathis, A., Britten, C.D., et al. (2015). A phase Ib dose-escalation study of the oral pan-PI3K inhibitor buparlisib (BKM120) in combination with the oral MEK1/2 inhibitor trametinib (GSK1120212) in patients with selected advanced solid tumors. Clin Cancer Res 21, 730–738.

Bhagwat, S.V., Gokhale, P.C., Crew, A.P., Cooke, A., Yao, Y., Mantis, C., Kahler, J., Workman, J., Bittner, M., Dudkin, L., et al. (2011). Preclinical characterization of OSI-027, a potent and selective inhibitor of mTORC1 and mTORC2: distinct from rapamycin. Mol Cancer Ther 10, 1394–1406.

Boison, D. (2013). Adenosine kinase: exploitation for therapeutic gain. Pharmacol Rev 65, 906–943.

Campana, L.G., Galuppo, S., Valpione, S., Brunello, A., Ghiotto, C., Ongaro, A., and Rossi, C.R. (2014). Bleomycin electrochemotherapy in elderly metastatic breast cancer patients: clinical outcome and management considerations. J Cancer Res Clin Oncol 140, 1557–1565.

Chandler, C., Liu, T., Buckanovich, R., and Coffman, L.G. (2019). The double edge sword of fibrosis in cancer. Transl Res 209, 55–67.

Chang, Y., Park, H., Yang, H.J., Lee, S., Lee, K.Y., Kim, T.S., Jung, J., and Shin, J.M. (2018). Cancer Drug Response Profile scan (CDRscan): A Deep Learning Model That Predicts Drug Effectiveness from Cancer Genomic Signature. Sci Rep 8, 8857.

Chen, K.G., and Sikic, B.I. (2012). Molecular pathways: regulation and therapeutic implications of multidrug resistance. Clin Cancer Res 18, 1863–1869.

Chia, S.K., Bramwell, V.H., Tu, D., Shepherd, L.E., Jiang, S., Vickery, T., Mardis, E., Leung, S., Ung, K., Pritchard, K.I., et al. (2012). A 50-gene intrinsic subtype classifier for prognosis and prediction of benefit from adjuvant tamoxifen. Clin Cancer Res 18, 4465–4472.

Chiu, Y.C., Chen, H.H., Zhang, T., Zhang, S., Gorthi, A., Wang, L.J., Huang, Y., and Chen, Y. (2019). Predicting drug response of tumors from integrated genomic profiles by deep neural networks. BMC Med Genomics 12, 18.

Colpitts, T.L., Billing, P., Granados, E., Hayden, M., Hodges, S., Roberts, L., Russell, J., Friedman, P., and Stroupe, S. (2002). Identification and immunohistochemical characterization of a mucin-like glycoprotein expressed in early stage breast carcinoma. Tumour Biol 23, 263–278.

Conley, S.J., Bosco, E.E., Tice, D.A., Hollingsworth, R.E., Herbst, R., and Xiao, Z. (2016). HER2 drives Mucin-like 1 to control proliferation in breast cancer cells. Oncogene 35, 4225–4234.

Corcoran, R.B., Atreya, C.E., Falchook, G.S., Kwak, E.L., Ryan, D.P., Bendell, J.C., Hamid, O., Messersmith, W.A., Daud, A., Kurzrock, R., et al. (2015). Combined BRAF and MEK Inhibition With Dabrafenib and Trametinib in BRAF V600-Mutant Colorectal Cancer. J Clin Oncol 33, 4023–4031.

Day, R.M., Yang, Y.Z., Suzuki, Y.J., Stevens, J., Pathi, R., Perlmutter, A., Fanburg, B.L., and Lanzillo, J.J. (2001). Bleomycin upregulates gene expression of angiotensin-converting enzyme via mitogen-activated protein kinase and early growth response 1 transcription factor. Am J Resp Cell Mol 25, 613–619.

Dellinger, R.W., Chen, G., Blevins-Primeau, A.S., Krzeminski, J., Amin, S., and Lazarus, P. (2007). Glucuronidation of PhIP and N-OH-PhIP by UDP-glucuronosyltransferase 1A10. Carcinogenesis 28, 2412–2418.

Dellinger, R.W., Fang, J.L., Chen, G., Weinberg, R., and Lazarus, P. (2006). Importance of UDP-glucuronosyltransferase 1A10 (UGT1A10) in the detoxification of polycyclic aromatic hydrocarbons: decreased glucuronidative activity of the UGT1A10139Lys isoform. Drug Metab Dispos 34, 943–949.

Dhillon, A.S., Hagan, S., Rath, O., and Kolch, W. (2007). MAP kinase signalling pathways in cancer. Oncogene 26, 3279–3290.

Gao, Y., and Liao, P. (2019). TRPM4 channel and cancer. Cancer Lettb 454, 66–69.

Garnett, M.J., Edelman, E.J., Heidorn, S.J., Greenman, C.D., Dastur, A., Lau, K.W., Greninger, P., Thompson, I.R., Luo, X., Soares, J., et al. (2012). Systematic identification of genomic markers of drug sensitivity in cancer cells. Nature 483, 570–575.

Gaspar, N., Sharp, S.Y., Pacey, S., Jones, C., Walton, M., Vassal, G., Eccles, S., Pearson, A., and Workman, P. (2009). Acquired resistance to 17-allylamino-17-demethoxygeldanamycin (17-AAG, tanespimycin) in glioblastoma cells. Cancer Res 69, 1966–1975.

Geeleher, P., Cox, N.J., and Huang, R.S. (2014). Clinical drug response can be predicted using baseline gene expression levels and in vitro drug sensitivity in cell lines. Genome Biol 15, R47.

Geeleher, P., Zhang, Z., Wang, F., Gruener, R.F., Nath, A., Morrison, G., Bhutra, S., Grossman, R.L., and Huang, R.S. (2017). Discovering novel pharmacogenomic biomarkers by imputing drug response in cancer patients from large genomics studies. Genome Res 27, 1743–1751.

Ghandi, M., Huang, F.W., Jane-Valbuena, J., Kryukov, G.V., Lo, C.C., McDonald, E.R., 3rd, Barretina, J., Gelfand, E.T., Bielski, C.M., Li, H., et al. (2019). Next-generation characterization of the Cancer Cell Line Encyclopedia. Nature 569, 503–508.

Grob, J.J., Amonkar, M.M., Karaszewska, B., Schachter, J., Dummer, R., Mackiewicz, A., Stroyakovskiy, D., Drucis, K., Grange, F., Chiarion-Sileni, V., et al. (2015). Comparison of dabrafenib and trametinib combination therapy with vemurafenib monotherapy on health-related quality of life in patients with unresectable or metastatic cutaneous BRAF Val600-mutation-positive melanoma (COMBI-v): results of a phase 3, open-label, randomised trial. Lancet Oncol 16, 1389–1398.

Gross, I., Bassit, B., Benezra, M., and Licht, J.D. (2001). Mammalian sprouty proteins inhibit cell growth and differentiation by preventing ras activation. J Biol Chem 276, 46460–46468.

Guan, N.N., Zhao, Y., Wang, C.C., Li, J.Q., Chen, X., and Piao, X. (2019). Anticancer Drug Response Prediction in Cell Lines Using Weighted Graph Regularized Matrix Factorization. Mol Ther Nucleic Acids 17, 164–174.

Gudernova, I., Vesela, I., Balek, L., Buchtova, M., Dosedelova, H., Kunova, M., Pivnicka, J., Jelinkova, I., Roubalova, L., Kozubik, A., et al. (2016). Multikinase activity of fibroblast growth factor receptor (FGFR) inhibitors SU5402, PD173074, AZD1480, AZD4547 and BGJ398 compromises the use of small chemicals targeting FGFR catalytic activity for therapy of short–stature syndromes. Hum Mol Genet 25, 9–23.

Guvenc Paltun, B., Mamitsuka, H., and Kaski, S. (2019). Improving drug response prediction by integrating multiple data sources: matrix factorization, kernel and network-based approaches. Brief Bioinform.

Hodges, L.M., Markova, S.M., Chinn, L.W., Gow, J.M., Kroetz, D.L., Klein, T.E., and Altman, R.B. (2011). Very important pharmacogene summary: ABCB1 (MDR1, P-glycoprotein). Pharmacogenet Genomics 21, 152–161.

Iorio, F., Knijnenburg, T.A., Vis, D.J., Bignell, G.R., Menden, M.P., Schubert, M., Aben, N., Goncalves, E., Barthorpe, S., Lightfoot, H., et al. (2016). A Landscape of Pharmacogenomic Interactions in Cancer. Cell 166, 740–754.

Kim, J.K., Ryll, R., Ishizuka, Y., and Kato, S. (2000). Identification of cDNAs encoding two novel nuclear proteins, IMUP-1 and IMUP-2, upregulated in SV40-immortalized human fibroblasts. Gene 257, 327–334.

Kim, S.J., An, H.J., Kim, H.J., Jungs, H.M., Lee, S., Ko, J.J., Kim, I.H., Sakuragi, N., and Kim, J.K. (2008). Imup-1 and imup-2 overexpression in endometrial carcinoma in Korean and Japanese populations. Anticancer Res 28, 865–871.

Li, L., Weinberg, C.R., Darden, T.A., and Pedersen, L.G. (2001). Gene selection for sample classification based on gene expression data: study of sensitivity to choice of parameters of the GA/KNN method. Bioinformatics 17, 1131–1142.

Li, P., Xiao, H.D., Xu, J., Ong, F.S., Kwon, M., Roman, J., Gal, A., Bernstein, K.E., and Fuchs, S. (2010). Angiotensin-converting enzyme N-terminal inactivation alleviates bleomycin-induced lung injury. Am J Pathol 177, 1113–1121.

Long, G.V., Stroyakovskiy, D., Gogas, H., Levchenko, E., de Braud, F., Larkin, J., Garbe, C., Jouary, T., Hauschild, A., Grob, J.J., et al. (2015). Dabrafenib and trametinib versus dabrafenib and placebo for Val600 BRAF-mutant melanoma: a multicentre, double-blind, phase 3 randomised controlled trial. Lancet 386, 444–451.

Lugowska, I., Kosela-Paterczyk, H., Kozak, K., and Rutkowski, P. (2015). Trametinib: a MEK inhibitor for management of metastatic melanoma. Onco Targets Ther 8, 2251–2259.

Masoumi-Moghaddam, S., Amini, A., and Morris, D.L. (2014). The developing story of Sprouty and cancer. Cancer Metastasis Rev 33, 695–720.

Merrill, G.F., Kurth, E.J., Hardie, D.G., and Winder, W.W. (1997). AICA riboside increases AMP-activated protein kinase, fatty acid oxidation, and glucose uptake in rat muscle. Am J Physiol 273, E1107–1112.

Miksicek, R.J., Myal, Y., Watson, P.H., Walker, C., Murphy, L.C., and Leygue, E. (2002). Identification of a novel breast- and salivary gland-specific, mucin-like gene strongly expressed in normal and tumor human mammary epithelium. Cancer Res 62, 2736–2740.

Nguyen, L., Dang, C.C., and Ballester, P.J. (2016). Systematic assessment of multi-gene predictors of pan-cancer cell line sensitivity to drugs exploiting gene expression data. F1000Res 5.

Paramanov, V., Tyurin, O., Polenkov, S., and Goldfarb, P.M. (2007). A safety and efficacy study of bleomycin sulfate and electroporation in patients with metastatic or locally recurrent breast cancer. Breast Cancer Research 9, SP5.

Pietanza, M.C., Waqar, S.N., Krug, L.M., Dowlati, A., Hann, C.L., Chiappori, A., Owonikoko, T.K., Woo, K.M., Cardnell, R.J., Fujimoto, J., et al. (2018). Randomized, Double-Blind, Phase II Study of Temozolomide in Combination With Either Veliparib or Placebo in Patients With Relapsed-Sensitive or Refractory Small-Cell Lung Cancer. J Clin Oncol 36, 2386–2394.

Rajapakse, V.N., Luna, A., Yamade, M., Loman, L., Varma, S., Sunshine, M., Iorio, F., Sousa, F.G., Elloumi, F., Aladjem, M.I., et al. (2018). CellMinerCDB for Integrative Cross-Database Genomics and Pharmacogenomics Analyses of Cancer Cell Lines. iScience 10, 247–264.

Rees, M.G., Seashore-Ludlow, B., Cheah, J.H., Adams, D.J., Price, E.V., Gill, S., Javaid, S., Coletti, M.E., Jones, V.L., Bodycombe, N.E., et al. (2016). Correlating chemical sensitivity and basal gene expression reveals mechanism of action. Nat Chem Biol 12, 109–116.

Reinhold, W.C., Sunshine, M., Liu, H., Varma, S., Kohn, K.W., Morris, J., Doroshow, J., and Pommier, Y. (2012). CellMiner: a web-based suite of genomic and pharmacologic tools to explore transcript and drug patterns in the NCI-60 cell line set. Cancer Res 72, 3499–3511.

Reinhold, W.C., Varma, S., Rajapakse, V.N., Luna, A., Sousa, F.G., Kohn, K.W., and Pommier, Y.G. (2015). Using drug response data to identify molecular effectors, and molecular “omic” data to identify candidate drugs in cancer. Hum Genet 134, 3–11.

Reinhold, W.C., Varma, S., Sunshine, M., Elloumi, F., Ofori-Atta, K., Lee, S., Trepel, J.B., Meltzer, P.S., Doroshow, J.H., and Pommier, Y. (2019). RNA Sequencing of the NCI-60: Integration into CellMiner and CellMiner CDB. Cancer Res 79, 3514–3524.

Robert, C., Grob, J.J., Stroyakovskiy, D., Karaszewska, B., Hauschild, A., Levchenko, E., Chiarion Sileni, V., Schachter, J., Garbe, C., Bondarenko, I., et al. (2019). Five-Year Outcomes with Dabrafenib plus Trametinib in Metastatic Melanoma. N Engl J Med 381, 626–636.

Robert, C., Karaszewska, B., Schachter, J., Rutkowski, P., Mackiewicz, A., Stroiakovski, D., Lichinitser, M., Dummer, R., Grange, F., Mortier, L., et al. (2015). Improved overall survival in melanoma with combined dabrafenib and trametinib. N Engl J Med 372, 30–39.

Robey, R.W., Pluchino, K.M., Hall, M.D., Fojo, A.T., Bates, S.E., and Gottesman, M.M. (2018). Revisiting the role of ABC transporters in multidrug-resistant cancer. Nat Rev Cancer 18, 452–464.

Rosenthal, T., and Gavras, I. (2009). Angiotensin inhibition and malignancies: a review. J Hum Hypertens 23, 623–635.

Ryoo, Z.Y., Jung, B.K., Lee, S.R., Kim, M.O., Kim, S.H., Kim, H.J., Ahn, J.Y., Lee, T.H., Cho, Y.H., Park, J.H., et al. (2006). Neoplastic transformation and tumorigenesis associated with overexpression of IMUP-1 and IMUP-2 genes in cultured NIH/3T3 mouse fibroblasts. Biochem Biophys Res Commun 349, 995–1002.

Sagredo, A.I., Sagredo, E.A., Pola, V., Echeverria, C., Andaur, R., Michea, L., Stutzin, A., Simon, F., Marcelain, K., and Armisen, R. (2019). TRPM4 channel is involved in regulating epithelial to mesenchymal transition, migration, and invasion of prostate cancer cell lines. J Cell Physiol 234, 2037–2050.

Seashore-Ludlow, B., Rees, M.G., Cheah, J.H., Cokol, M., Price, E.V., Coletti, M.E., Jones, V., Bodycombe, N.E., Soule, C.K., Gould, J., et al. (2015). Harnessing Connectivity in a Large-Scale Small-Molecule Sensitivity Dataset. Cancer Discov 5, 1210–1223.

Sotiriou, C., and Pusztai, L. (2009). Gene-expression signatures in breast cancer. N Engl J Med 360, 790–800.

Suphavilai, C., Bertrand, D., and Nagarajan, N. (2018). Predicting Cancer Drug Response using a Recommender System. Bioinformatics 34, 3907–3914.

Uchiyama, S., Itoh, H., Naganuma, S., Nagaike, K., Fukushima, T., Tanaka, H., Hamasuna, R., Chijiiwa, K., and Kataoka, H. (2007). Enhanced expression of hepatocyte growth factor activator inhibitor type 2-related small peptide at the invasive front of colon cancers. Gut 56, 215–226.

Vaidyanathan, A., Sawers, L., Gannon, A.L., Chakravarty, P., Scott, A.L., Bray, S.E., Ferguson, M.J., and Smith, G. (2016). ABCB1 (MDR1) induction defines a common resistance mechanism in paclitaxel- and olaparib-resistant ovarian cancer cells. Br J Cancer 115, 431–441.

Valladares-Ayerbes, M., Iglesias-Diaz, P., Diaz-Prado, S., Ayude, D., Medina, V., Haz, M., Reboredo, M., Antolin, S., Calvo, L., and Anton-Aparicio, L.M. (2009). Diagnostic accuracy of small breast epithelial mucin mRNA as a marker for bone marrow micrometastasis in breast cancer: a pilot study. J Cancer Res Clin Oncol 135, 1185–1195.

Vassilev, L.T., Vu, B.T., Graves, B., Carvajal, D., Podlaski, F., Filipovic, Z., Kong, N., Kammlott, U., Lukacs, C., Klein, C., et al. (2004). In vivo activation of the p53 pathway by small-molecule antagonists of MDM2. Science 303, 844–848.

Wallden, B., Storhoff, J., Nielsen, T., Dowidar, N., Schaper, C., Ferree, S., Liu, S., Leung, S., Geiss, G., Snider, J., et al. (2015). Development and verification of the PAM50-based Prosigna breast cancer gene signature assay. BMC Med Genomics 8, 54.

Wei, D., Liu, C., Zheng, X., and Li, Y. (2019). Comprehensive anticancer drug response prediction based on a simple cell line-drug complex network model. BMC Bioinformatics 20, 44.

Xu, Q.S., and Liang, Y.Z. (2001). Monte Carlo cross validation. Chemometr Intell Lab 56, 1–11.

Yang, W., Soares, J., Greninger, P., Edelman, E.J., Lightfoot, H., Forbes, S., Bindal, N., Beare, D., Smith, J.A., Thompson, I.R., et al. (2013). Genomics of Drug Sensitivity in Cancer (GDSC): a resource for therapeutic biomarker discovery in cancer cells. Nucleic Acids Res 41, D955–961.

Zoppoli, G., Regairaz, M., Leo, E., Reinhold, W.C., Varma, S., Ballestrero, A., Doroshow, J.H., and Pommier, Y. (2012). Putative DNA/RNA helicase Schlafen-11 (SLFN11) sensitizes cancer cells to DNA-damaging agents. Proc Natl Acad Sci U S A 109, 15030–15035.

