## Supplemental Figure S1, S2, S3, S4, S5 and S6 for "Predicting Tumor Response to Drugs based on Gene-Expression Biomarkers of Sensitivity Learned from Cancer Cell Lines"

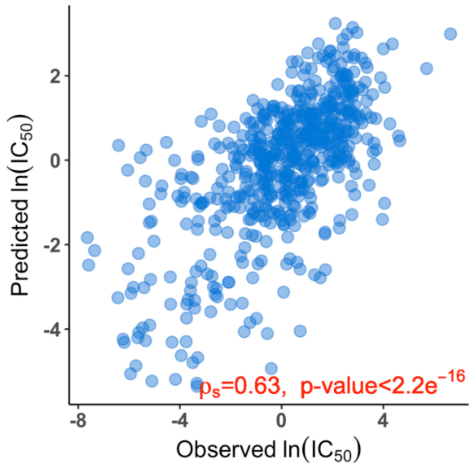


**Figure S1**. Scatter plot of predicted and observed ln($IC_{50}$) values for trametinib in the 571 cancer cell lines with both gene expression data and $IC_{50}$ data for trametinib.

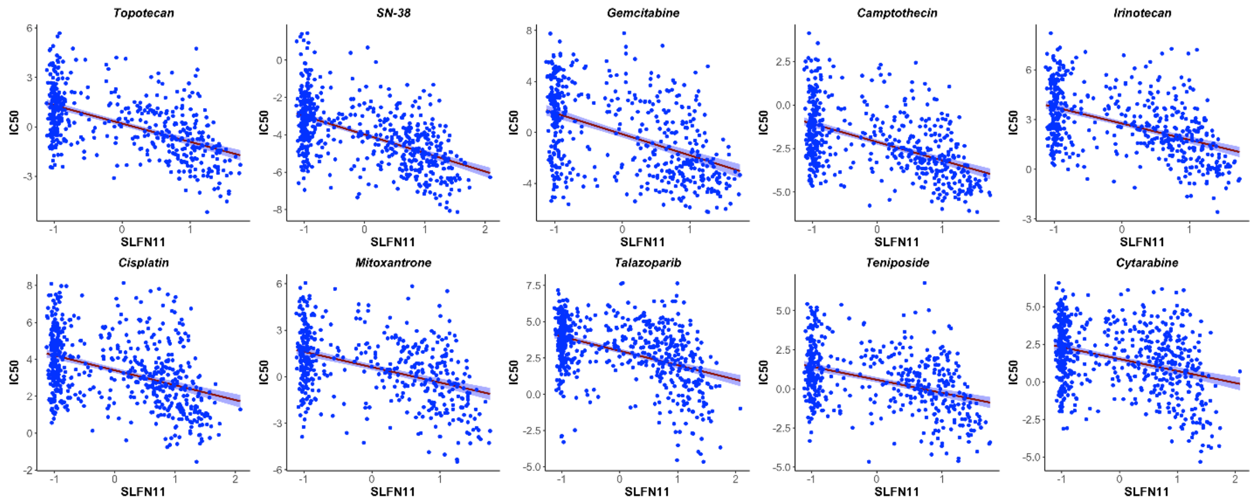


**Figure S2**. Inverse correlation between *SLFN11* expression (Z score) in cancer cell lines and the observed ln($IC_{50}$) values of the 10 drugs, suggesting that higher *SLFN11* expression in cancer cell lines is positively correlated with the sensitivity of the cell lines to those drugs

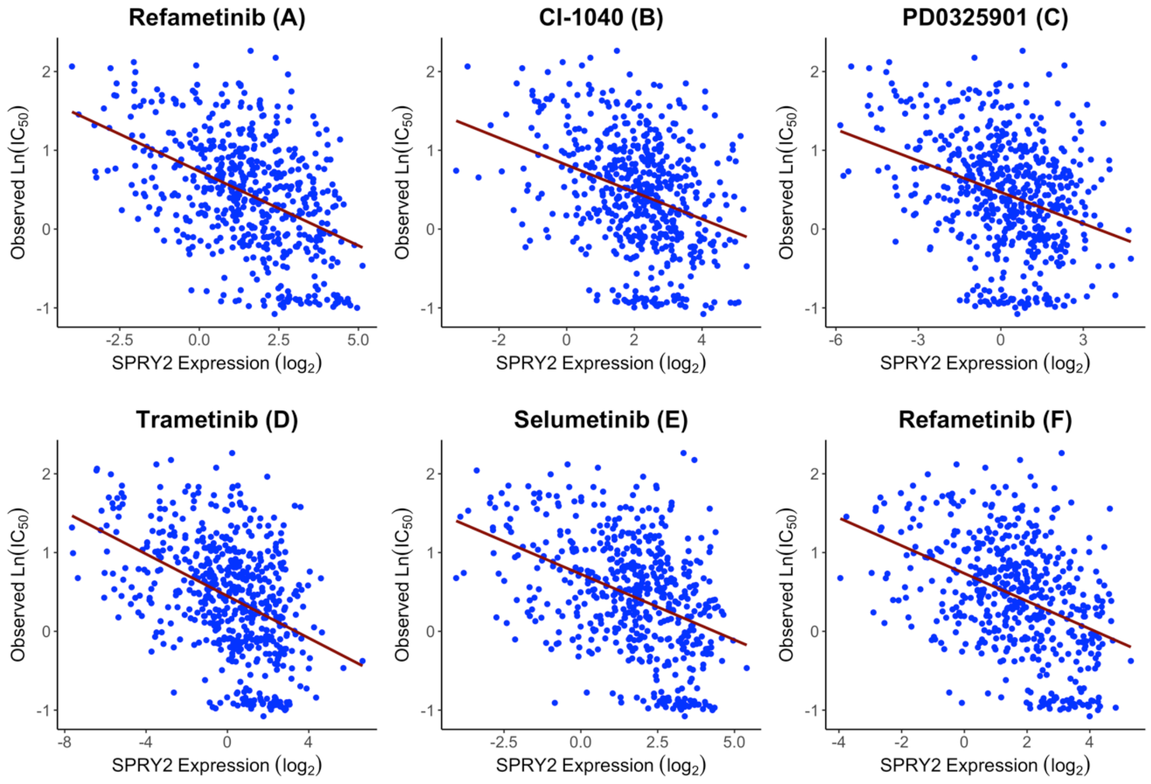


**Figure S3**. Inverse correlation between *SPRY2* expression (Z score) in cancer cell lines and the observed $\ln(IC_{50})$ of the six MEK inhibitors for those cell lines. Higher *SPRY2* expression was associated with lower $\ln(IC_{50})$ value (more sensitive to these drugs).

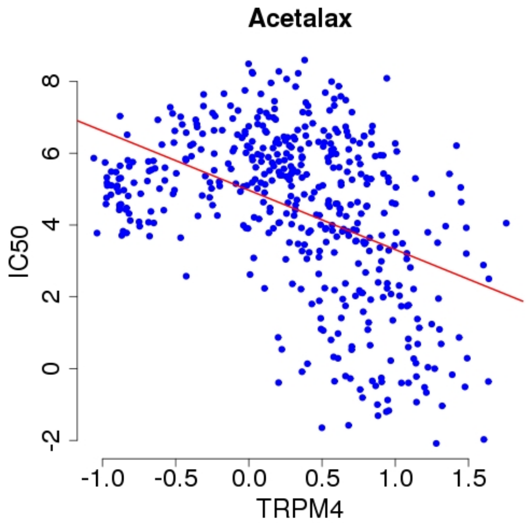


**Figure S4**. *TRPM4* expression (Z score) in cancer cell lines is inversely correlated with observed $\ln(IC_{50})$ values of acetalax for the cell lines, indicating high *TRPM4* expression is associated with high sensitivity (low $IC_{50}$) to acetalax.

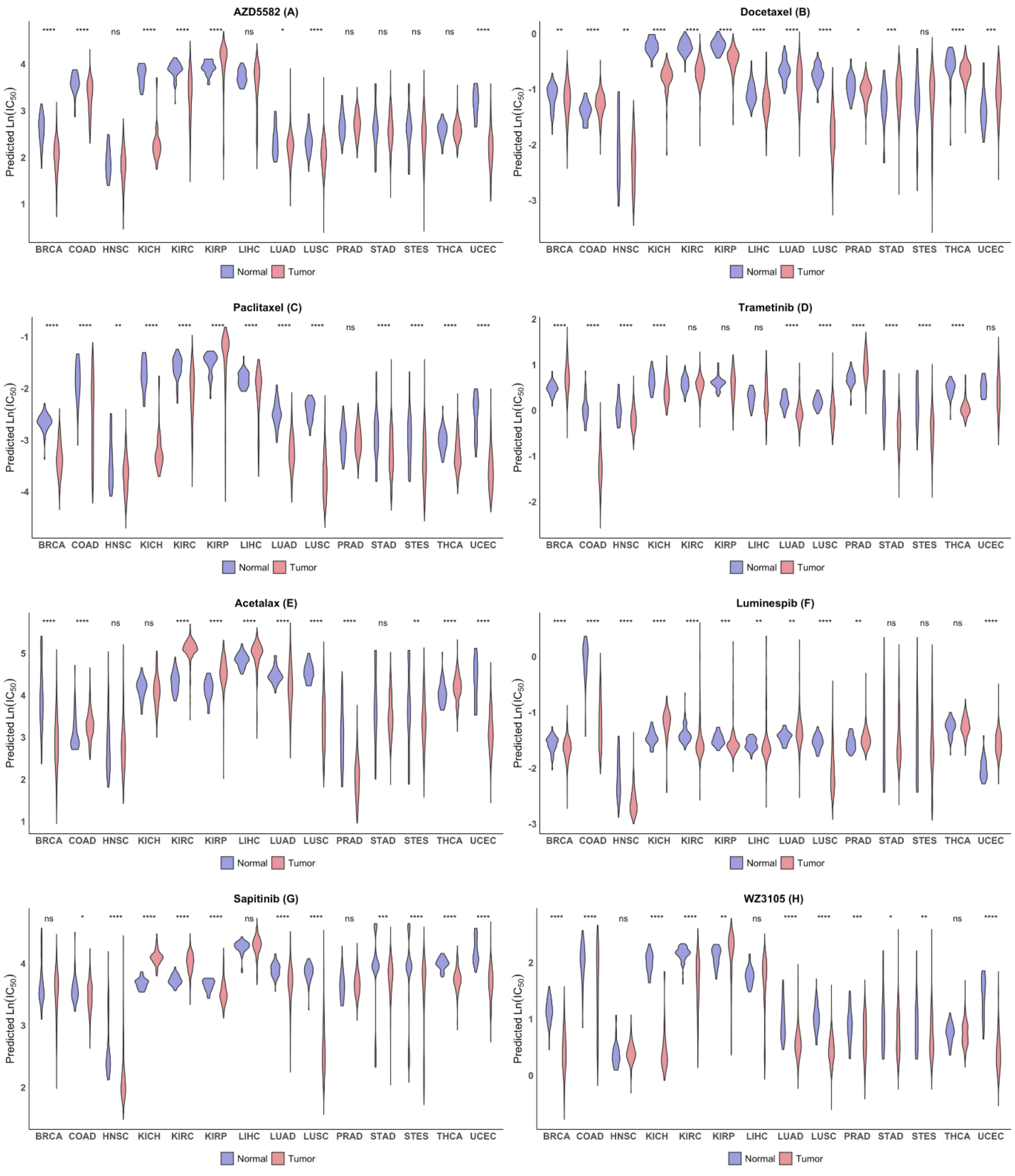


**Figure S5.** Drugs that are predicted to have high tumor-to-normal sensitivity for some tumor types. Violin plots of predicted ln($IC_{50}$) values in tumor (pink) and normal (blue) tissue for the eight drugs that showed the ratio of tumor-to-normal sensitivity exceeding 2.7 (1 logarithmic unit) for at least one of 14 tissue types. The ln($IC_{50}$ ) values of the drugs were predicted based on the RNA-seq data of the tumor and normal tissue samples from TCGA. Violin plots for normal and tumor samples from the same tissue type are shown as side-by-side pairs with their TCGA type on the x-axis. See Figure 4 legend for additional description of the violin plots. Statistical significance based on a two-tailed Mann-Whitney-Wilcoxon rank sum test is shown above each pair with ns: p > 0.05, *: p <= 0.05, **: p <= 0.01, ***: p <= 0.001, ****: p <= 0.0001.

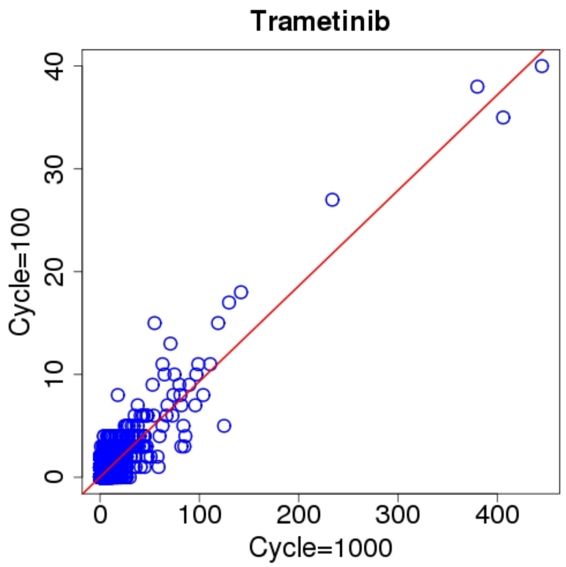


**Figure S6**. Scatter plot of the counts of genes selected into the sets of 30 chromosomes from two independent runs with 100 runs and 1,000 runs, respectively.
