## Supplemental Table S1, S2, S6, S9, S10 and S11 for "Predicting Tumor Response to Drugs based on Gene-Expression Biomarkers of Sensitivity Learned from Cancer Cell Lines"

**Table S1**. Summaries statistics of correlations between the observed and predicted $IC_{50}$ of the 453 drugs in the test set

| Correlation | Min. | 1st Qu. | Median | Mean | 3rd Qu. | Max. |
|---|---|---|---|---|---|---|
| $\rho_P$ | -0.1990 | 0.3710 | 0.4660 | 0.4573 | 0.5510 | 0.7660 |
| $\rho_S$ | -0.1800 | 0.3580 | 0.4370 | 0.4267 | 0.5070 | 0.6800 |

**Table S2**. The 10 most predictable drugs

| GDSC ID | Name | Target | Testing $\rho_p$ | Testing $\rho_s$ |
|---|---|---|---|---|
| 1047 | Nutlin-3a(-) | MDM2 | 0.729 | 0.680 |
| 1909 | Venetoclax | BCL2 | 0.766 | 0.568 |
| 1003 | Camptothecin | TOP1 | 0.654 | 0.664 |
| 1088 | Irinotecan | TOP1 | 0.663 | 0.647 |
| 1190 | Gemcitabine | DNA synthesis | 0.645 | 0.644 |
| 1372 | Trametinib | MEK1, MEK2 | 0.653 | 0.635 |
| 1814 | Nelarabine | DNA synthesis | 0.728 | 0.547 |
| 1563 | EPZ5676 | DOT1L | 0.647 | 0.629 |
| 252 | WZ3105 | NTRK and SRC kinase | 0.650 | 0.620 |
| 1931 | MIRA-1 | TP53 | 0.667 | 0.601 |

**Table S6**. TCGA Tumor types

| Tumor type | TCGA Code |
|---|---|
| adrenocortical carcinoma | ACC |
| bladder urothelial carcinoma | BLCA |
| breast invasive carcinoma | BRCA |
| cervical squamous cell carcinoma and endocervical adenocarcinoma | CESC |
| cholangiocarcinoma | CHOL |
| colon adenocarcinoma | COAD |
| lymphoid neoplasm diffuse large B-cell lymphoma | DLBC |
| glioblastoma multiforme | GBM |
| head and neck squamous cell carcinoma | HNSC |
| kidney chromophobe | KICH |
| kidney renal clear cell carcinoma | KIRC |
| kidney renal papillary cell carcinoma | KIRP |
| acute myeloid leukemia | LAML |
| brain lower grade glioma | LGG |
| liver hepatocellular carcinoma | LIHC |
| lung adenocarcinoma | LUAD |
| lung squamous cell carcinoma | LUSC |
| mesothelioma | MESO |
| ovarian serous cystadenocarcinoma | OV |
| pancreatic adenocarcinoma | PAAD |
| pheochromocytoma and paraganglioma | PCPG |
| prostate adenocarcinoma | PRAD |
| rectum adenocarcinoma | READ |

| | |
|---|---|
| sarcoma | SARC |
| skin cutaneous melanoma | SKCM |
| stomach adenocarcinoma | STAD |
| Esophagus-Stomach cancers | STES |
| testicular germ cell tumors | TGCT |
| thyroid carcinoma | THCA |
| thymoma | THYM |
| uterine corpus endometrial carcinoma | UCEC |
| uterine carcinosarcoma | UCS |
| uveal melanoma | UVM |

**Table S9**. Main parameters of the GA/KNN algorithm used for the analyses of all datasets

| Parameter | Value |
|---|---|
| population size | 5,000 |
| maximum number of generations | 2,000 |
| chromosome length (*d*) | 30 |
| number of nearest neighbors (*k*) | 3 |
| number of independent GA/KNN runs | 100 |

**Table 10.** Training and testing performances for various combinations of *k* and *d*

| KNN (*k*) | Chromosome length (*d*) | Training | | Testing | |
|---|---|---|---|---|---|
| | | Pearson | Spearman | Pearson | Spearman |
| 1 | 20 | 0.926 | 0.911 | 0.624 | 0.611 |
| | 30 | 0.931 | 0.917 | 0.605 | 0.567 |
| | 40 | 0.933 | 0.920 | 0.621 | 0.586 |
| 3 | 20 | 0.899 | 0.871 | 0.634 | 0.615 |
| | 30 | 0.906 | 0.880 | 0.638 | 0.624 |
| | 40 | 0.912 | 0.890 | 0.620 | 0.589 |
| 5 | 20 | 0.880 | 0.849 | 0.634 | 0.618 |
| | 30 | 0.890 | 0.864 | 0.625 | 0.606 |
| | 40 | 0.898 | 0.873 | 0.638 | 0.612 |

**Table S11**. Comparison of training and testing performances between two independent runs with 100 and 1,000 runs, respectively.

| Run | Training | | Testing | |
|---|---|---|---|---|
| | Pearson | Spearman | Pearson | Spearman |
| 100 Runs | 0.908 | 0.886 | 0.627 | 0.608 |
| 1,000 Runs | 0.913 | 0.890 | 0.658 | 0.637 |
